# Pro-survival roles for p21(Cip1/Waf1) in Non-Small Cell Lung Cancer

**DOI:** 10.1101/2024.05.21.595102

**Authors:** SJ Cutty, FA Hughes, P Ortega-Prieto, S Desai, P Thomas, LV Fets, M Secrier, AR Barr

## Abstract

Quiescence is a reversible state of proliferative arrest, distinct from senescence. While cancer is a disease of dysregulated proliferation, cancer cells can retain the ability to enter quiescence which confers advantages to tumour cells by protecting them from chemotherapy or by allowing metastasis to distant sites. Multiple mechanisms exist to induce and maintain quiescence that are not yet fully understood. Here, we show that high expression of the CDK inhibitor p21^Cip1/Waf1^ correlates with a poor prognosis in *TP53* wild-type, but not *TP53* mutant, non-small cell lung cancer (NSCLC) patients. Using quantitative single-cell imaging of genetically-engineered NSCLC reporter cell lines, we show that *TP53* wild-type NSCLC cells can enter a p21-dependent spontaneous quiescent state, downstream of replication stress. Furthermore, p21 expression confers survival advantages to *TP53* wild-type NSCLC cells, both under normal proliferation and in response to chemotherapy. We also show that p21 can promote tumour relapse by allowing cells to recover from both G1 and G2 arrest states after drug removal. Together, our data suggest that targeting p21 function in *TP53* wild-type tumours could lead to better outcomes for chemotherapy treatment in NSCLC patients.

**Statement of Significance:** We show that *TP53*WT Non-Small Cell Lung Cancer cells can enter a p21-dependent spontaneous quiescent state and that p21 maintains the viability of NSCLC cells, is chemoprotective and can promote tumour relapse.

## Introduction

Lung cancer is the deadliest cancer worldwide and non-small cell lung cancer (NSCLC) comprises 85% of cases (1,2). The five-year survival rate for NSCLC is a dismal 15.9% (3), thus there is a desperate need for new therapeutic strategies. Approximately 80% of patients present with inoperable disease and are essentially incurable. Even where patients initially respond to therapy, they frequently relapse with resistant disease. Therefore, it is essential to identify mechanisms driving inherent and acquired resistance to chemotherapy, to reduce rates of local and distant relapse and increase patient survival.

Tumour relapse can be driven by dormant, or quiescent, cancer cells reinitiating proliferation (4). Quiescence (or G0) is a reversible cell cycle arrest, where cells temporarily exit proliferative cycles for either long or short periods of time (5). By their nature, quiescent cells are not targeted by standard cytotoxic chemotherapy aimed at killing proliferating cells. Cycles of chemotherapy aim to capture quiescent cells as they move between proliferating and arrested states. However, depending on the time that cells reside in quiescence and the fraction of quiescent cells within a tumour, chemotherapy cycles may be insufficient to completely eradicate cancer cells. To complicate matters further, quiescence is not a single cellular state but a set of poorly defined states (5–7), currently defined by a lack of proliferative markers rather than markers specific to quiescent subtypes. To reduce the incidence of tumour relapse, we need better characterisation of quiescent subtypes within tumours and new treatment strategies to either eradicate quiescent cancer cells or prevent them from re-entering proliferative cycles.

One quiescent subtype recently identified is induced in response to intrinsic DNA damage, or replication stress (8–11). This type of quiescence is dependent on p53-p21^Cip1/Waf1^ (p21) signalling (8,12,13). p21 is a cyclin-dependent kinase (CDK) inhibitor and a transcriptional target of p53 (14,15). During cell cycle entry, CyclinD:CDK4/6 and CyclinE:CDK2 cooperate to promote hyperphosphorylation and inactivation of the transcriptional repressor protein pRb (16). pRb hyperphosphorylation leads to the release and activation of a family of transcriptional activators, E2F1-3, which drive the transcription of multiple genes required for cell cycle entry (17). In response to intrinsic DNA damage, or replication stress, generated during normal DNA replication in S-phase, p53 is stabilised and drives p21 expression, such that p21 protein starts to accumulate during the G2 phase of the cell cycle. At these low levels of DNA damage, p21 rarely reaches a threshold that is capable of inhibiting CDK activity in G2 cells (18). Therefore, if the DNA damage is not repaired, then both the DNA damage and p21 protein are inherited by the daughter cells, where p21 expression increases further, inhibits CDK activity, pRb remains active and cells enter quiescence (8,19–22). Since cells in p21-dependent quiescence do eventually re-enter proliferative cycles this arrest is distinct from senescence (9). Knocking out p21 (p21KO) in cells generates increased levels of basal DNA damage (8). These data suggest that p21-dependent quiescence provides time for cells to repair, or prepare to repair, DNA damage, before resuming proliferative cycles, to maintain genome stability.

As a CDK inhibitor, p21 is most simply thought of as a tumour suppressor gene, preventing cells with DNA damage from proliferating and propagating mutations. p21KO mouse models are consistent with this, since mice developed spontaneous tumours, albeit with a longer latency than p53KO mice (23). p21KO mice are also more susceptible to tumorigenesis induced by carcinogens (24–26). However, *CDKN1A*, the gene encoding p21, is rarely mutated in human cancers (27). Moreover, p21 has been described to have pro-tumorigenic properties (28,29). For example, clinical studies in patients with advanced solid tumours (30) and in rectal carcinoma patients (31) indicated that low p21 expression correlates with improved disease-free survival. More mechanistic studies suggest that p53-independent upregulation of p21 may lead to dysregulated p21 expression which could drive genomic instability by disrupting DNA replication and increasing levels of replication stress (32). Whether p21 plays a tumour suppressive or pro-tumorigenic role in any particular tumour is, therefore, likely to depend on cellular context.

In NSCLC, p21 expression is heterogeneous but is frequently higher within tumour cells than in surrounding normal tissue (33). Data from the Human Protein Atlas (www.proteinatlas.org; (34)) suggests that high p21 expression correlates with a poor prognosis in lung cancer patients (Figure 1A). Since wild-type (WT) *TP53* is retained in approximately 50% of NSCLCs, these data, together with previous observations made by us and others (18–22), lead us to hypothesise that *TP53*WT NSCLC cells may have pro-tumorigenic or pro-survival properties that could be linked to the ability to enter p21-dependent quiescence. We reasoned, that by allowing NSCLC cells to temporarily exit proliferative cycles, p21-dependent quiescence may provide time for NSCLC cells to repair DNA damage and protect them from chemotherapy (Figure 1B).

**Figure 1.**
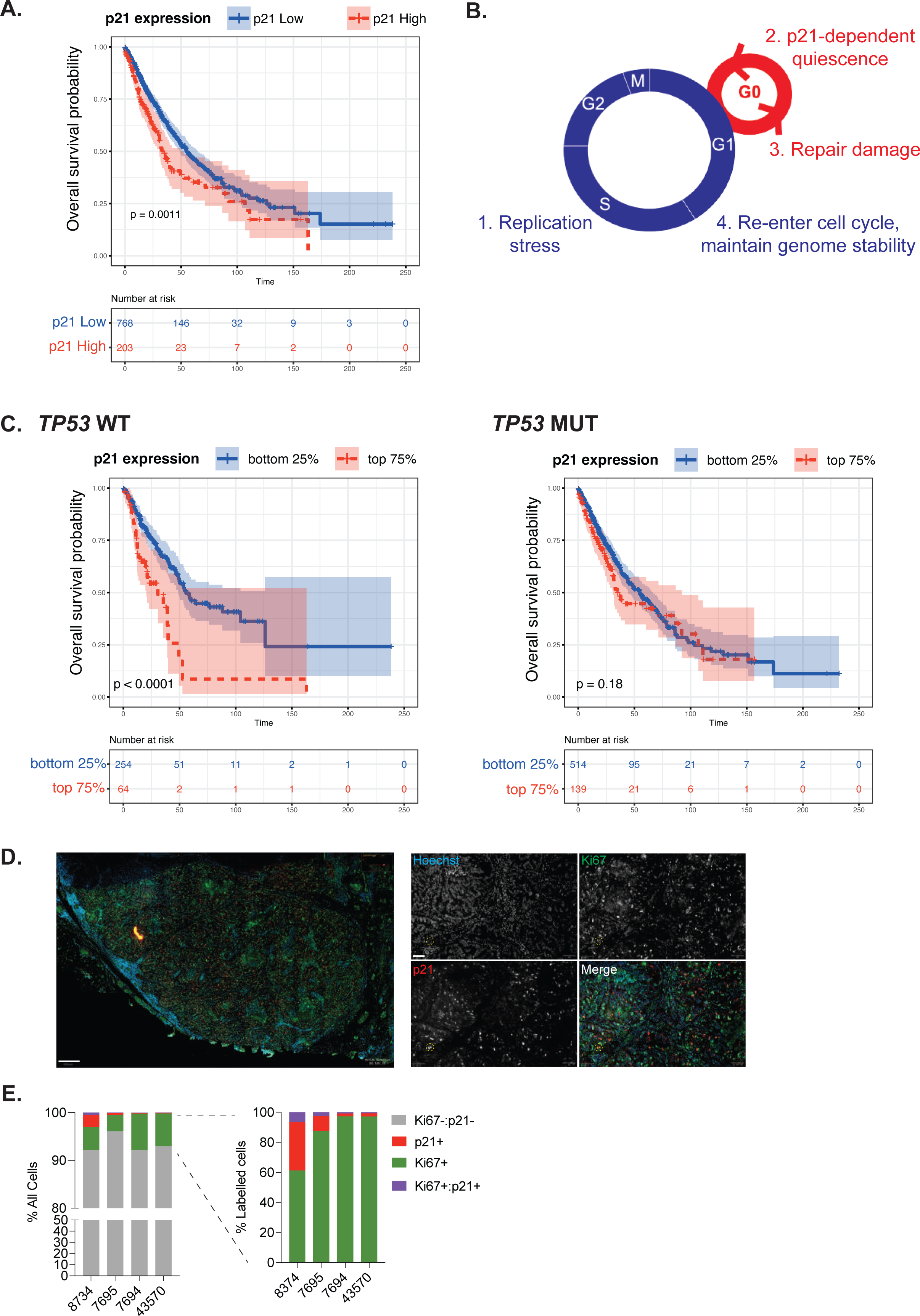
High p21 expression correlates with poor prognosis in *TP53*WT NSCLC. **A.** Graph replotted from data in the Human Protein Atlas. P21 high and low groups are defined based in the 67.52 optimised expression cut-off specified on the Human Protein Atlas website. **B.** Hypothetical model for beneficial role of p21 for NSCLC growth and druvival. **C.** TCGA analyses showing overall survival probability separated by *TP53* status and level of p21 (*CDKN1A*) mRNA expression. **D.** Image of *TP53*WT human lung tumour (patient 8734) immunostained for Hoechst to label nuclei (blue), Ki67 (red) and p21 (green in merged images). On the left is a zoomed-out tumour section (scale bar 400 μm). On the right are zoomed-in sections (scale bar 100 μm). An example of Ki67+/p21+ cells is highlighted with a yellow circle. **E.** Quantification of p21+, Ki67+ and Ki67+/p21+ cells from human tumours. On the left are all Hoechst-positive tumour cells. On the right, the same data are re-plotted with just those cells that were labelled for either p21 or Ki67.

In this study, we aimed to investigate the role(s) of p21 in NSCLC. We find that high p21 expression correlates with a poor prognosis specifically in *TP53*WT, and not *TP53*mutant, NSCLC. Using quantitative, single-cell imaging we show that p21-dependent quiescence exists in *TP53*WT NSCLC and is pro-survival in proliferating NSCLC cells. *TP53*WT NSCLC cells lacking p21 are more sensitive to chemotherapy, not because the p21-dependent quiescent state is chemoprotective but because cells lacking p21 can no longer sustain prolonged G1 or G2 arrest states. Post-chemotherapy treatment, we show that a fraction of cells residing in these p21-dependent G1- and G2-arrested states are quiescent, and not senescent, since these cells can reawaken and drive proliferation once more. Therefore, our data suggest that p21-dependent quiescence could drive tumour relapse in *TP53*WT NSCLC cells and that targeting p21 in combination with chemotherapy could improve patient outcomes.

## Results

### High levels of p21 correlate with a poor prognosis in *TP53* wild-type NSCLC

p21 has been shown to act as both a tumour suppressor and an oncogene, depending on context (27). Here, we wanted to investigate the potential pro-tumorigenic, or pro-survival, properties of p21 in NSCLC since a high level of p21 protein correlates with a worse prognosis in this disease (Figure 1A) and we hypothesised that p21 may provide a fitness advantage to NSCLC cells by allowing tumour cells with replication stress that persists beyond S-phase, to enter a p21-dependent quiescent state (G0) after mitosis (Figure 1B; (18,19,21,22)).

If our hypothesis is correct, we would expect to see that p21 expression correlates with a worse prognosis only in *TP53*WT tumours, where cells can respond to replication stress through p53-dependent p21 expression (18), and not in *TP53*mutant tumours. Therefore, we analysed NSCLC cases in The Cancer Genome Atlas (TCGA), separated by *TP53* status and level of *CDKN1A* (encoding p21) expression. Consistent with our hypothesis, a high level of *CDKN1A* expression only correlated with a poor prognosis in *TP53*WT NSCLC and not in *TP53*mutant tumours (Figure 1C). We observed the same correlation when analysing protein expression data (Supplementary Figure 1).

p21 expression has been reported to be higher in tumours than in surrounding normal tissue (35). However, p21 is frequently used as a marker of senescence in tissues and tumours. We wanted to identify if p21-expressing cells in lung tumours maintain proliferative capacity and whether some of these cells could be quiescent and not senescent. Since there are no good markers that uniquely distinguish quiescent from senescent cells, we opted to use Ki67 staining as a marker of proliferative potential. It was recently shown that Ki67 is a graded marker of cellular proliferation, that is expressed in quiescent cells but that decreases in protein level the longer cells are held in quiescence (36). We immunostained *TP53*WT NSCLC tumours for p21 and Ki67 and quantified the fraction of p21+, Ki67+ and p21+/Ki67+ cells (Figure 1D). We identified a small and variable fraction of p21+/Ki67+ cells, indicating the possibility of p21-dependent quiescent cells in patient tumours (Figure 1E). Cells negative for both p21 and Ki67 are most likely terminally-differentiated cells.

In summary, our analyses show that high p21 expression correlates with poor NSCLC patient prognosis specifically in *TP53*WT tumours, which account for approximately 50% of NSCLC cases. We provide evidence that p21 expressing cells in *TP53*WT tumours can be proliferative and are not terminally arrested in senescence.

### *TP53* wild-type NSCLC cells can enter a p21-dependent quiescent state

We first investigated whether proliferating NSCLC cells could enter a “spontaneous” quiescent state and if this was p21-dependent. Spontaneous here refers to cells entering quiescence in the presence of full growth media and in the absence of contact inhibition (37). We selected a panel of NSCLC cell lines – five *TP53*WT (NCI-H460, A549, NCI-H1944, NCI-H1666, NCI-H1563), three *TP53*mutant (NCI-H1299, CORL23, NCI-H358) and one that has a mutation causing psi-p53 to be expressed (NCI-H1650 (38)). We validated that *TP53*WT cells had an intact p53-p21 pathway by treating cells with the Mdm2 inhibitor, Nutlin-3, and quantifying p21 expression by single-cell imaging. All *TP53*WT and none of the *TP53*mutant cell lines induced p21 expression upon Nutlin-3 treatment (Figure S2A). psi-p53 has been suggested to be transcriptionally inactive(39). However, we observed upregulation of p21 protein expression in NCI-H1650 cells in response to Nutlin-3 and downregulation of p21 after p53 depletion, more similar to *TP53*WT than *TP53*mutant cells (Figure S2A,B). Therefore, we consider the NCI-H1650 cell line as an intermediate between *TP53*WT and *TP53*mutant cells that retains the ability to regulate p21 in a p53-dependent manner.

We developed a high-throughput single-cell imaging assay to identify quiescent cells. We plated cells at low cell density, then added EdU, a nucleoside analogue incorporated into cells undergoing DNA replication, for the final 24h before fixing and staining cells. In addition to EdU labelling, we immunostained cells for hyperphosphorylated pRb (P-Rb), a marker for cells that have passed the Restriction Point and committed to proliferation (40,41) (Figure 2A,B; Figure S3A,B). EdU negative are cells have not undergone S-phase for at least 24h and can be said to be quiescent. Similarly, cells that are negative for hyperphosphorylated Rb can be considered quiescent. Using this assay, we identified a fraction of spontaneously quiescent cells across all cell lines tested, ranging from 1-47% of quiescent cells (EdU assay) and 10-66% quiescent cells (P-Rb assay; Figure 2C, Supplementary Figure 3C). The fraction of quiescent cells identified in the two assays differs due to the different assay timescales but the two measurements correlate (Supplementary Figure 3D).

**Figure 2.**
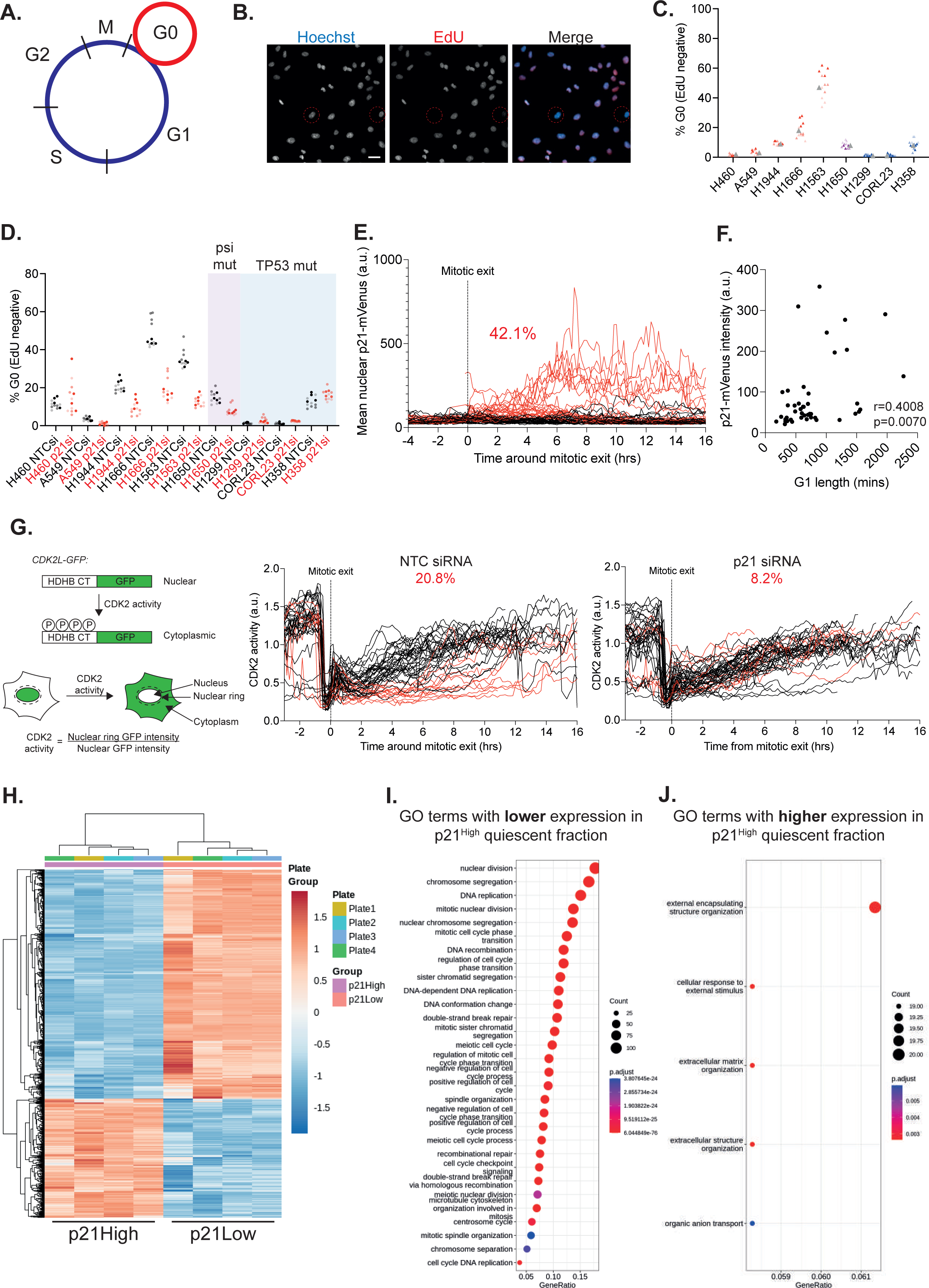
p53 wild-type NSCLC cells can enter a p21-dependent quiescent state. **A.** Schematic shows assay principle – EdU is taken up by cells that are actively proliferating during the 24h incubation (blue circle). Cells in quiescence (G0, red circle) will not be labelled with EdU. **B.** Sample images of A549 cells labelled with EdU to quantify the quiescent fraction. Red circles represent EdU negative, quiescent cells. EdU is in red and Hoechst in blue in merged image. Scale bar 20 μm. **C.** Graph shows quantification of percentage of EdU negative (G0) cells across cell lines. Data are plotted as superplots, n=3, 4 technical replicates per experiment. Grey triangle represents the mean. *TP53*WT cell lines are shown in red, psi-p53 mutant is shown in purple and *TP53*mutant cell lines in blue. **D.** Graph shows the effect of p21 depletion by siRNA on the fraction of quiescent cells within each cell line, measured using the EdU assay. Data are plotted as superplots, n=3, 4 technical replicates per experiment. Non-targeting control (NTC) siRNA values are shown in black, p21 siRNA in red. **E.** Quantification of nuclear p21-mVenus in single NCI-H1944 cells over time. All single cell traces are aligned to mitotic exit. Red curves represent cells that exit mitosis with high p21-mVenus levels (defined as > 60 a.u. at 5h post-mitotic exit). Percentage in red is the fraction of cells that enter a p21^High^ state (proxy for quiescence, 24/57 cells (42.1%). **F.** Correlation between G1 length and p21-mVenus levels. **G.** Left: schematic to show how the CDK2 activity sensor works (44). Right: quantification of CDK2 activity in single A549 cells over time. All single cell traces are aligned to mitotic exit. Red curves represent cells that exit mitosis with low CDK2 activity (defined as < 0.5 at 6h post-mitotic exit). Percentages in red are the fraction of cells in each condition that enter a CDK2^Low^ activity state (proxy for quiescence). NTC siRNA, 14/48 cells (20.8%). p21 siRNA, 4/49 cells (8.2%). **H.** Heatmap shows clustering of significantly differentially expressed genes between p21-High versus p21-Low cells based on RNA-seq profiling. Four repeats per condition were profiled. Red represents increased expression, blue represents decreased expression. **I.** GO analysis of genes with lower expression in p21^High^ cells. **J.** GO analysis of genes with higher expression in p21^Low^ cells.

To determine if entry into this spontaneous quiescent state is p21-dependent in NSCLC cells, we depleted p21 using siRNA (Supplementary Figure 3E). Four of the five *TP53*WT cell lines and the psi-p53 cell line all showed a reduction in the quiescent fraction when p21 was depleted (Figure 2D, Supplementary Figure S3F). None of the three *TP53*mutant lines showed any change in the quiescent fraction after p21 depletion. These data show that p21-dependent quiescence exists in *TP53*WT NSCLC cells.

Quiescence is a reversible cell cycle arrest state and cells enter and exit quiescence on different timescales. While our fixed cell assay is useful for characterising quiescence across cell lines in a high-throughput manner, we need to quantify quiescence in a more dynamic manner. To do this, we generated fluorescent reporter *TP53*WT NSCLC cell lines to quantify entry and exit into quiescence. First, we generated an NCI-H1944 cell line where we labelled endogenous PCNA with mRuby to follow cell cycle dynamics and to accurately define phase transitions (42) and tagged endogenous p21 with an mVenus fluorophore (43) to follow p21 expression (Supplementary Figure S3G-J). Live cell imaging revealed that p21 expression is heterogeneous, highest in G1 cells and that high p21 expression correlates with entry into quiescence. Approximately 42% of NCI-H1944 cells exit mitosis with high p21 levels and remain in a period of quiescence (longer G1 phase) before re-entering the cell cycle (Figure 2E, red curves), while the remaining cells have low levels of p21 and re-enter S-phase more rapidly (Figure 2E, black curves, 2F; Supplementary Movie 1; (18,20)). We also generated an A549 cell line expressing mRuby-PCNA and a CDK2 activity sensor (44) (Figure 2G, Supplementary Figure 3K). In cells that enter quiescence, we anticipate CDK2 activity to decrease after mitotic exit, as p21 levels increase (44,45). Indeed, we observed that approximately 20% of A549 cells downregulate CDK2 activity upon exiting mitosis and that this is p21-dependent (red curves, Figure 2G).

We wanted to determine what, in addition to p21 expression, defined these spontaneously quiescent NSCLC cells. As well as demonstrating localisation changes throughout the cell cycle, PCNA also displays expression changes throughout the cell cycle, with G0/G1 cells displaying the lowest PCNA protein levels. We used FACS to separate p21-mVenus^High^ mRuby-PCNA^Low^ (‘p21^High^’) quiescent cells from p21-mVenus^Low^ mRuby-PCNA^High^ (‘p21^Low^’) proliferating cells (Supplementary Figure 3L) and validated the two populations by immunostaining for additional cell cycle markers (Supplementary Figure 3M). We characterised the transcriptomes of these two populations by RNAseq (Figure 2H; Supplementary Figure 3N). As expected, genes involved in regulating the cell cycle were significantly reduced in p21^High^ quiescent cells (Figure 2I; Table S1), further validating this as a quiescent population. Genes that were significantly enriched in p21^High^ quiescent cells included genes encoding proteins involved in cell interactions with the external environment (Figure 2J; Table S2). These included genes associated with GO terms involving response to external stimuli (IL1B, TNFSF14, NR1H4, PHEX, MAP1LC3C, WIPI1, SCX, FOLR1, CASP1, RRAGD, TLR5, PDK4, CD68, CDKN1A, AGT, ZFYVE1, BMT2, NAMPT, GABARAPL1), in extracellular matrix and structure organisation (TGM3, ADAMTS14, ADAMTS7, IL6, SCX, COL4A3, COL17A1, LRP1, ADAMTSL4, COL5A1, MMP11, PBXIP1, MMP2, ADAMTS13, LOXL2, AGT, LUM, ADAMTS10, LAMB3, BMP1) and in organic anion transport (IL1B, KMO, SLC22A11, CEACAM1, SLC16A4, SLC6A12, SLC4A10, FOLR1, SLC26A9, CES1, SLC4A3, SLC23A1, SLC27A1, SLC17A5, AGT, ABCC6, PTGES, PSAP, CYB5R1). The enrichment of mRNAs for ADAMTS (a disintegrin and metalloproteinase with thrombospondin motifs) and MMP (matrix metalloproteinase) suggest that p21^High^ quiescent cells may be more invasive than their proliferative counterparts.

Together, these data show that p21-dependent quiescence exists in *TP53*WT, but not *TP53*mutant, NSCLC. p21-High quiescent cells have unique transcriptional profiles suggesting they are a discrete cell state. There is considerable heterogeneity amongst cell lines in the fraction of cells in p21-dependent quiescence at any one time and within cell lines in the length of time spent in quiescence by individual cells. For the remainder of this work, we focus our efforts on trying to understand the causes and consequences of p21-dependent quiescence in NSCLC. Therefore, from here, we focus only on *TP53*WT NSCLC cells.

### Replication stress can drive entry into p21-dependent quiescence

We, and others, have previously shown that a key cause of p21 upregulation and entry into p21-dependent quiescence is intrinsic DNA damage caused by DNA replication stress (18,19,21,22). Therefore, we wanted to see if the spontaneously quiescent cells identified in *TP53*WT NSCLC had persistent DNA damage. We pulse-labelled cells with EdU 24 hours prior to fixation and immunostained for p21 and 53BP1, the latter of which forms nuclear bodies around damaged DNA and is associated with persistent replication stress (Figure 3A; (46)). We identified EdU-negative or p21-High quiescent cells and quantified the fraction of these cells with at least one 53BP1 nuclear body, compared to EdU positive and p21-Low proliferating cells (Figure 3B). We consistently observed across *TP53*WT NSCLC cell lines that a higher fraction of EdU negative and p21-High cells had at least one 53BP1 nuclear body, more than is observed in proliferating cells in the same population (Figure 3C, Supplementary Figure 4A). This suggests that entry into the quiescent state in *TP53*WT NSCLC is linked to persistent DNA damage.

**Figure 3.**
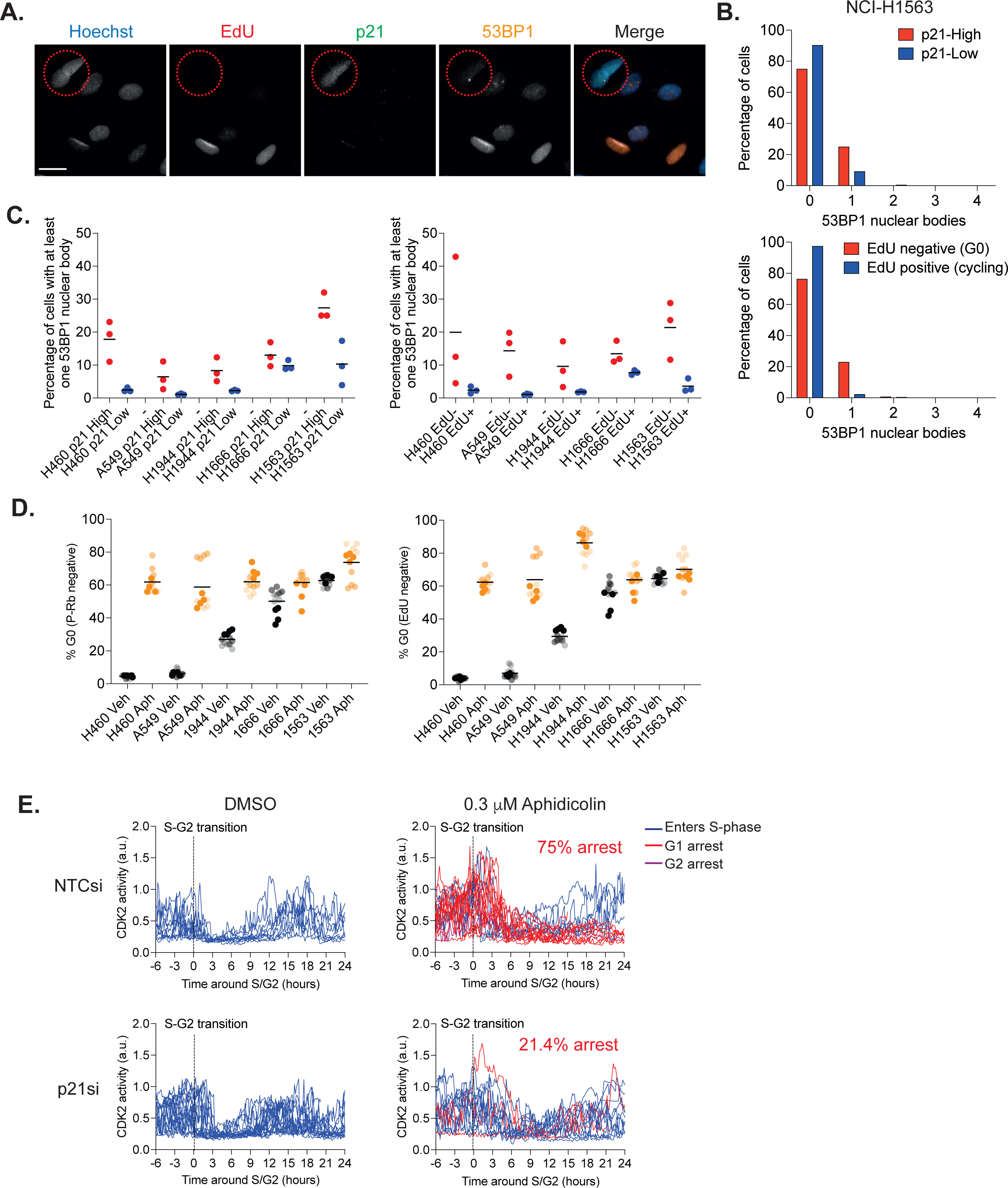
Replication stress can drive entry into p21-dependent quiescence. **A.** Images of NCI-H1563 cells labelled with EdU for 24h and stained for p21 and 53BP1. Red circle highlights an EdU negative (quiescent) cell with a 53BP1 nuclear body and expressing p21. EdU is red, p21 is green, 53BP1 is orange and Hoechst is blue in merged image. Scale bar is 10 μm. **B.** Graphs show the percentage of p21-High (upper panel) or EdU negative (lower panel) quiescent cells with a 53BP1 nuclear body, compared to proliferating cells. Data are representative of n=3. **C.** Summary of percentage of quiescent cells (red) with at least one 53BP1 nuclear body, compared to proliferating cells (blue). Black line represents the mean of n=3. **D.** Percentage of cells arresting in quiescence after vehicle (grey dots) or low-dose (0.3 μM) aphidicolin (orange dots) treatment. Data are plotted as superplots, n=3, 4 technical replicates per experiment. Black line is the mean of n=3. **E.** Quantification of CDK2 activity in single A549 mRuby-PCNA CDK2L-GFP cells after DMSO or 0.3 μM aphidicolin treatment. All single cell traces are aligned to the S-G2 transition. Blue curves represent cells that re-enter S-phase after DMSO or aphidicolin treatment, red curves represent cells that arrest in G1 and purple curves (just one cell in NTCsi + aphidicolin) represent cells arresting in G2. Percentages in red are the fraction of cells that enter an arrest state after aphidicolin treatment. NTC siRNA, 15/20 cells (75%). p21 siRNA, 3/14 cells (21.4%).

To test if replication stress can push cells into a quiescent state in our *TP53*WT NSCLC cell lines, we treated cells with a low dose of the DNA polymerase-α inhibitor, aphidicolin, to induce more replication stress. We verified that this dose of aphidicolin induced replication stress in these cell lines by quantifying the number of 53BP1 bodies per cell and the level of p21 expression, both of which increase after aphidicolin treatment (Supplementary Figure 4B-D). We observed an increased fraction of cells arresting in quiescence after low-dose aphidicolin treatment (Figure 3D). We also performed live cell imaging with our A549 mRuby-PCNA CDK2L-GFP cell line, treated with either a negative control or p21 siRNA and with or without aphidicolin treatment to enable us to monitor entry into quiescence in real-time (Figure 3E). We observed that in the presence of p21 a higher fraction of A549 cells treated with aphidicolin enter a quiescent state post-mitosis (75% in NTCsi versus 21.4% in p21si).

Together our data suggest that replication stress correlates with a p21-dependent quiescent state and can drive entry into that state in *TP53*WT NSCLC cells.

### Loss of p21 leads to propagation of DNA damage into S-phase, perturbation of cell cycle dynamics and spontaneous cell death

Tumour cells often exhibit high levels of DNA damage in the form of replication stress as a consequence of oncogene activation (47,48). Therefore, we wanted to examine the consequences of the loss of p21 on *TP53*WT NSCLC cells on genome stability and proliferative potential.

We depleted p21 using siRNA in mRuby-PCNA expressing A549 and NCI-H1944 cells to track cell cycle phenotypes and cell fates by timelapse imaging. After acute p21 depletion, we observed a higher rate of spontaneous cell death (Figure 4A,B; Supplementary Figure 5A). Cell death was not restricted to a single cell cycle phase. To test the hypothesis that p21 helps to maintain cell viability in *TP53*WT NSCLC cells, we used CRISPR/Cas9 to disrupt the p21 gene (*CDKN1A*) in our five *TP53*WT NSCLC cell lines to generate p21 knockout (p21KO) cell lines. We were only able to generate p21KO clonal lines in three of the five cell lines (A549 mRuby-PCNA, NCI-H1944 mRuby-PCNA and NCI-H460; Supplementary Figure 5B,C), despite multiple targeting attempts. As expected, all p21KO cell lines had a reduced fraction of cells entering spontaneous quiescence (Supplementary Figure 5D,E). We used timelapse imaging of p21WT and p21KO A549 and NCI-H1944 mRuby-PCNA cells to track cell cycle phenotypes and cell fates. Again, we observed increased spontaneous cell death in p21KO cell lines compared to p21WT cells (Figure 4C). Of note, these rates of death were lower than those observed after acute depletion of p21 by siRNA (Figure 4B) which may reflect p21KO cell lines adapting to p21 loss.

**Figure 4.**
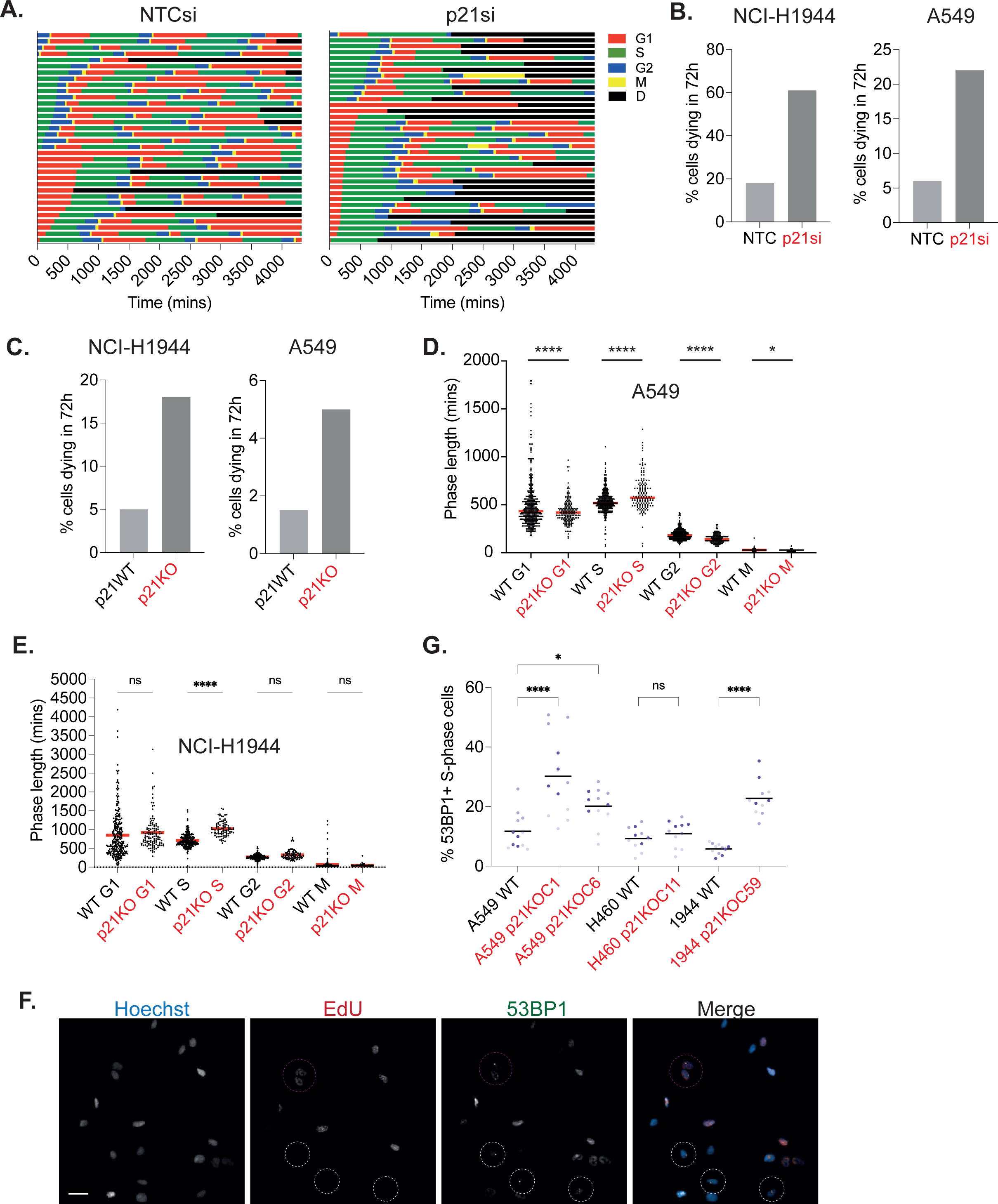
Loss of p21 leads to propagation of DNA damage into S-phase, perturbation of cell cycle dynamics and spontaneous cell death. **A.** Cell cycle phase plots for NCI-H1944 mRuby-PCNA cells transfected with NTC or p21-targeting siRNA. 34 cells for NTCsi and 36 cells for p21si. “D” stands for death. **B.** Percentage of cells that die during the 72h imaging period after acute depletion of p21 by siRNA for NCI-H1944 cells (left) and A549 cells (right). **C.** Percentage of cells that die during the 72h imaging period in p21KO cells in NCI-H1944 cells (left) and A549 cells (right). **D.** Graph shows measurement of cell cycle phase lengths in A549 p21WT and p21KO cells. Each dot is the phase length in an individual cell, the red line represents the mean. Statistical significance was calculated using an unpaired t-test, between each cell cycle phase, with Welch’s correction. *p<0.05, ****p<0.0001. **E.** Graph shows measurement of cell cycle phase length in NCI-H1944 p21WT and p21KO cells. Each dot is the phase length in an individual cell, the red line represents the mean. Statistical significance was calculated using an unpaired t-test, between each cell cycle phase, with Welch’s correction. ****p<0.0001, ns = not significant. **F.** Representative images of NCI-H1944 mRuby-PCNA p21KO cells pulse-labelled with EdU (red in merged image), immunostained for 53BP1 (green in merged image) and stained with Hoechst to label nuclei (blue in merged image). Purple circled cells are those where 53BP1 bodies are present in S-phase. White circled cells are 53BP1 bodies present outside S-phase. Scale bar 20 μm. **G.** Quantification of percentage of S-phase cells with at least one 53BP1 nuclear body present. Data are presented as superplots, n=3 repeats with 4 technical replicates. Black line represents the mean. Statistical significance was calculated using a one-way ANOVA, between each p21KO clone and the p21WT counterpart. ns = not significant, *p<0.05, ****p<0.0001.

We analysed cell cycle phase lengths to assess the impact of p21 loss. In A549 mRuby-PCNA cells, we observed a reduction in G1 length, largely due to the loss of cells entering the prolonged G0/G1-like state of quiescence, an increase in the length of S-phase, and a decrease in the lengths of G2 and mitosis (Figure 4D). We observed a similar increase in S-phase length in p21KO NCI-H1944 mRuby-PCNA cells compared to p21WT cells (Figure 4E). We hypothesised that with a reduced ability to enter quiescence, which would normally allow cells to repair any inherited DNA damage, more DNA damage may be propagated into S-phase and that this may contribute to a longer S-phase length. Therefore, we quantified the levels of DNA damage in p21WT and p21KO NSCLC cell lines. We saw an overall increase in the levels of nuclear γH2AX in p21KO versus p21WT cells, consistent with an overall increase in genomic instability in the absence of p21-dependent quiescence (Supplementary Figure 5F). To identify damage specifically in S-phase cells, we pulse-labelled cells with EdU for 30 mins before fixation and immunostained with 53BP1 (Figure 4F). We observed an increase in the fraction of S-phase cells with DNA damage in p21KO versus p21WT cells, that was statistically significant in two out of the three p21KO lines (Figure 4G). This suggests that more DNA damage is being propagated into S-phase cells in the absence of p21-dependent quiescence.

Our data suggest that p21-dependent quiescence is required for the maintenance of genome stability, limiting the amount of DNA damage being propagated into S-phase cells, and that p21 acts to maintain the proliferative potential of *TP53*WT NSCLC cells by promoting cell fitness.

### Loss of p21 sensitises *TP53*WT NSCLC cells to chemotherapy

Since quiescence can protect cancer cells from chemotherapy agents that target cycling cells, we wanted to see if loss of p21-dependent quiescence in *TP53*WT NSCLC cells would sensitise them to chemotherapy.

We performed dose curves in A549, NCI-H460 and NCI-H1944 cell lines to determine sensitivity to three chemotherapy agents used to treat NSCLC: cisplatin, gemcitabine and etoposide. We observed a small fraction of cells remaining in all cell lines, even at the highest doses of drug (grey boxes, Supplementary Figure 6A). The fraction of cells surviving drug treatment was positively correlated with the fraction of quiescent cells in each cell line (Figure 5A, Supplementary Figure 6B). To determine if had an impact on the response of these cell lines to chemotherapy, we compared survival of p21WT and p21KO cell lines treated with cisplatin, gemcitabine and etoposide. We observed that p21KO A549 and NCI-H1944 cells were significantly more sensitive to etoposide and gemcitabine than p21WT cells, while p21KO NCI-H460 cells were significantly more sensitive to cisplatin than p21WT cells (Figure 5B, Supplementary Figure 6C).

**Figure 5.**
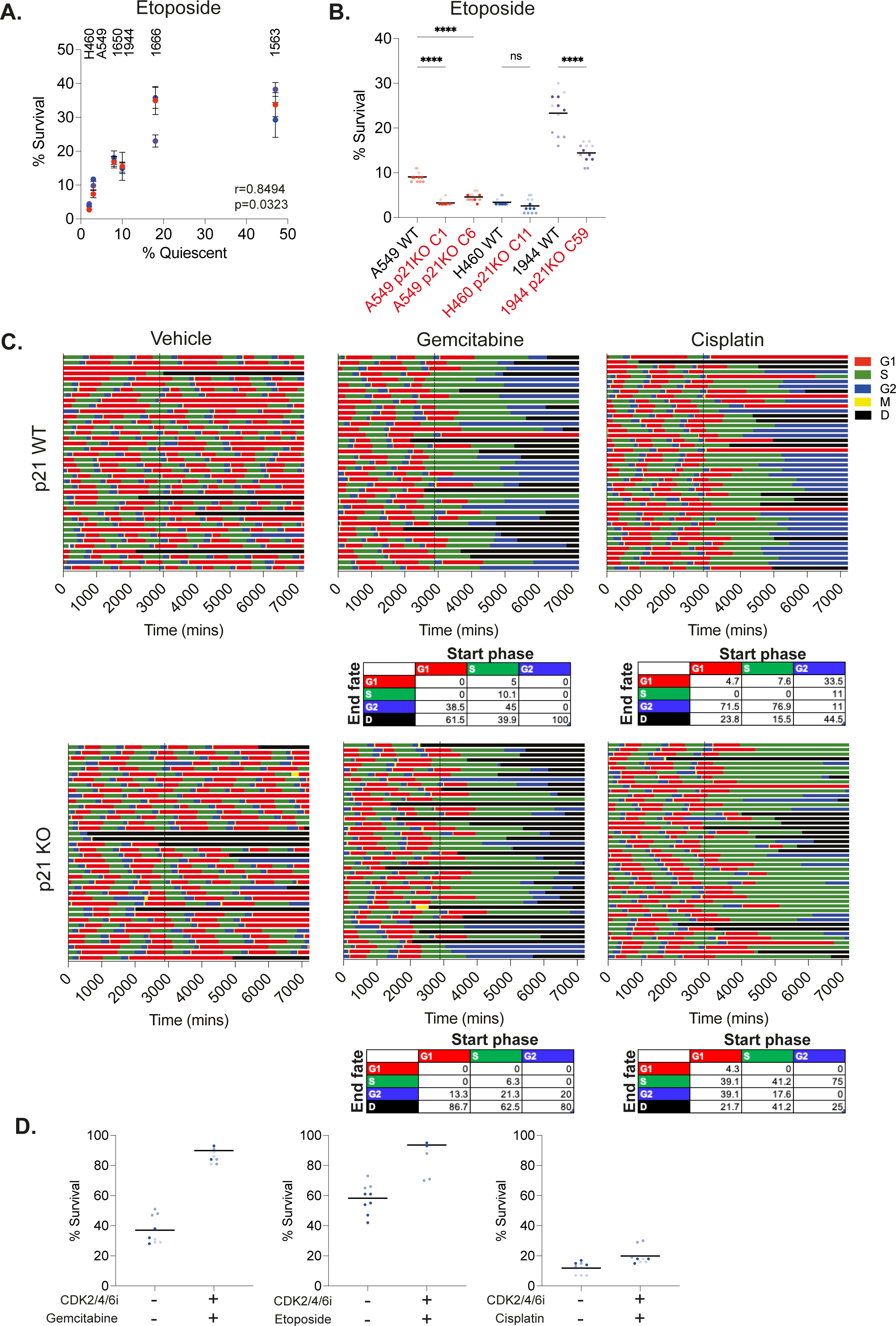
Loss of p21 sensitises *TP53*WT NSCLC cells to chemotherapy. **A.** Correlation between fraction of cells surviving etoposide treatment and quiescent fraction in that cell line. Cells were treated for three days with 5 μM etoposide. Mean +/- std for each repeat plotted and n=3 for each cell line. **B.** p21WT and p21KO cell lines were treated for three days with 5 μM of etoposide and surviving fraction was calculated. Data are plotted as superplots, n=3 biological repeats and four technical repeats per experiment. Black line represents the mean. One-way ANOVA was used to calculate statistical significance. **** = p<0.0001, ns = not significant. **C.** Cell cycle phase plots for A549 mRuby-PCNA p21WT (upper panels) and p21KO (lower panels) cells treated with gemcitabine or cisplatin for three days. Cells were imaged for two days before drug addition (marked by vertical black lines at 2880 mins). Tables underneath show the percentage of cells that started in each fate upon drug addition (G1, S or G2) and their end fate (G1, S, G2 or death (D)). **D.** Graphs showing percentage of cells surviving drug treatment (gemcitabine, etoposide or cisplatin) in quiescence. Cells were arrested in quiescence for one day in CDK2/4/6i prior to three days treatment with chemotherapy agent (still in the presence of CDK2/4/6i). Data are plotted as superplots, n=3 biological replicates with three technical replicates per repeat. Black line represents the mean.

To determine if it was the p21-dependent quiescent G0/G1 state that we observe in unperturbed cell populations that confers chemo-protection, or another function of p21, we used timelapse imaging. We imaged A549 and NCI-H1944 mRuby-PCNA expressing cells for two days to establish the cell cycle state before drug treatment. We then added either gemcitabine or cisplatin and continued to image for three days to track cell fates (Figure 5C, Supplementary Figure 6D). Cells that started in G1 upon drug treatment were equally likely to die as those cells in S or G2, suggesting residing in p21-dependent quiescence prior to drug treatment does not protect cells from chemotherapy. For gemcitabine, which targets DNA synthesis in S-phase, this is perhaps not too surprising and indeed, gemcitabine-treated cells that were in G1 at time of treatment complete S-phase before arresting in G2 or dying. However, for cisplatin, which forms DNA adducts and blocks DNA repair, and therefore can affect cells at all cell cycle stages, we found this more surprising. Most cisplatin-treated G1 cells arrest in G2 or die (Figure 5C; Supplementary Figure 6D), perhaps because cells in G1 do not mount a sufficient DNA damage response to cisplatin and so progress into the cell cycle. Specifically in A549 cells, we did observe that approximately one third of cells that were in G2 upon cisplatin addition survived in a p21-dependent G1 state and that cells in any state upon either gemcitabine or cisplatin treatment could enter a p21-dependent G2 arrest (Figure 5C).

We next asked if *TP53*WT NSCLC cells held in quiescence for the duration of chemotherapy treatments would be protected from chemotherapy. Addition of the CDK2/4/6 inhibitor, ebvaciclib (49), was able to arrest NCI-H1944 cells in quiescence (Supplementary Figure 6E). Therefore, we held NCI-H1944 cells in quiescence with CDK2/4/6 inhibitor and then treated arrested cells with gemcitabine, etoposide or cisplatin for three days. While cisplatin was able to kill quiescent NSCLC cells as efficiently as it killed proliferating cells, gemcitabine and etoposide were not (Figure 5D). These results are consistent with their mechanism of action and suggest that a sufficiently prolonged quiescence would not be sufficient to protect cells from cisplatin but may protect them from etoposide and gemcitabine.

Our data show that loss of p21 sensitises cells to chemotherapy. This is not due to a chemoprotective action of cells residing in a p21-dependent quiescent state during drug treatment being as this state may be too transient to confer protection but instead is due to p21 promoting cytoprotective prolonged G1 and G2 arrests.

### p21-dependent quiescence could drive tumour relapse

The prolonged p21-dependent G1 and G2 arrest states observed after chemotherapy treatment raise the question of how stable these arrest states are i.e. are these cells transiently arrested (quiescent) or terminally arrested (senescent)? These arrested cells express high levels of p21 and have enlarged nuclei, suggesting that these surviving cells could be in, or progressing towards, senescence (Figure 6A). However, these features are shared with quiescent cells (50). Therefore, we wanted to investigate if a population of p21-dependent quiescent cells were present post-drug treatment by determining if surviving cells can re-initiate cell proliferation, as a model of tumour relapse.

**Figure 6.**
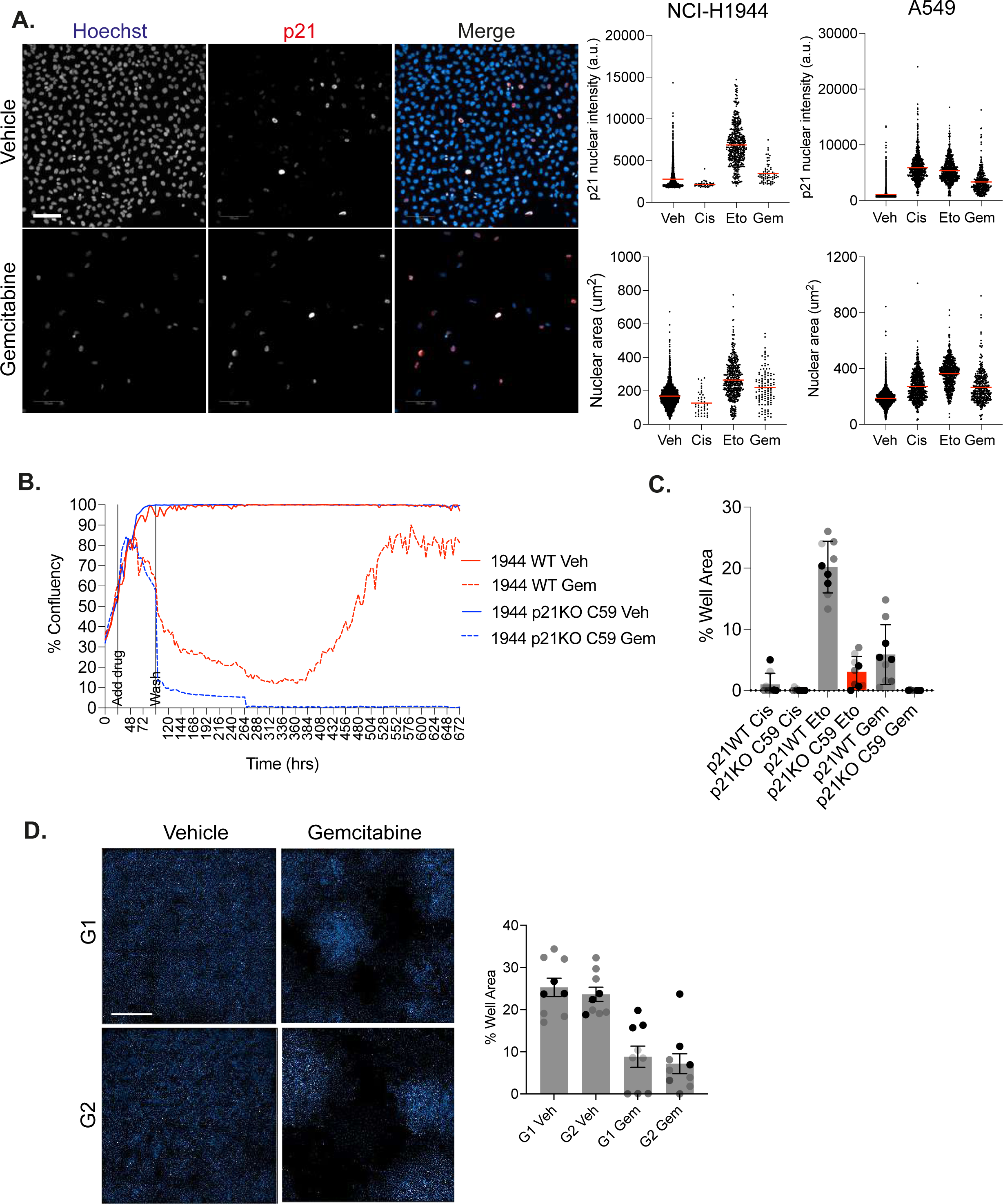
p21-dependent quiescence could drive tumour relapse. **A.** Representative images and quantification from A549 cells showing increase in p21 levels and nuclear area three days post-treatment with vehicle or gemcitabine. Hoechst is in blue and p21 is in red in merged images. Scale bar is 100μm. Graphs on right are single-cell quantifications of p21 nuclear intensity (upper panels) and nuclear area (lower panels) three days post-treatment with indicated drugs. Veh – vehicle, Cis - cisplatin, Eto – etoposide and Gem-gemcitabine. **B.** Representative graph of cell growth over time of NCI-H1944 p21WT (red curves) and p21KO cells (blue curves) treated with vehicle (solid line) or gemcitabine (dashed line) taken from one field of view. Cells were treated with drug for three days before the drug was washed out (‘wash’) and replaced with normal growth media to track any cell re-growth. **C.** Quantification of percentage of well covered by NCI-H1944 nuclei at the end of the relapse assay (28 days). Mean +/-stdev of n=3 are shown and individual wells are plotted for each technical repeat. **D.** Representative images of Hoechst stained NCI-H1944 nuclei from vehicle- or gemcitabine-treated G1 and G2 cells allowed to recover from drug treatment for two weeks. Scale bar is 1mm. Quantification of well area occupied by nuclei two weeks after sorting either G1 or G2 vehicle- or gemcitabine-treated cells and replating in drug-free media.

We treated p21WT and p21KO A549, NCI-H1944 and NCI-H460 cells for three days with cisplatin or gemcitabine before washing out the drug and imaging long-term cell growth. After the initial wave of drug-induced cell death and several days of no proliferation, p21WT cells started to proliferate again, a phenomenon that was much more rare in p21KO cells (Figure 6B,C). This was not a widespread event but individual colonies form and can eventually can recolonise the well (Figure 6C, Supplementary Figure 7A-C, Supplementary Movies 3-6). These data suggest that p21 can maintain a quiescent pool of cells after drug treatment that can drive population regrowth.

Since our single-cell imaging assays had identified that post-chemotherapy treatment, cells could arrest in G1 or, more frequently, G2 (Figure 5C, Supplementary Figure 6D), we wanted to ask if cells arresting in G1 or G2 were equally likely to re-enter the cell cycle after drug removal in p21WT cells. We also noticed that the colonies that regrew had different morphologies. Some colonies contained cells that were large and flat while other colonies contained cells that more closely resembled untreated cells (Supplementary Figure 7D, blue bounding edge) and wondered if these different cell morphologies represented whether cells had re-entered the cell cycle from G1 or G2. Therefore, we treated p21WT NCI-H1944 cells for three days with gemcitabine and then FACS sorted individual G1 or G2 cells into 96-well plates (Supplementary Figure 7E). We left cells for three weeks and assessed colony regrowth and cell morphology. We observed that cells were equally capable of re-entering the cell cycle from both G1 and G2 arrested populations and did not notice any difference in cell morphology between the two populations (Figure 6D).

Together, our data show that p21 can maintain a pool of quiescent cells post chemotherapy treatment that could drive tumour relapse.

## Discussion

We have shown how the CDK inhibitor, p21, can exert pro-survival properties in *TP53*WT NSCLC. Expression of p21, downstream of replication stress, can drive NSCLC cells into a quiescent state post-mitosis. In the absence of p21, DNA damage incurred through replication stress is propagated into the subsequent S-phase and *TP53*WT cells that have lost p21 have decreased viability. Moreover, p21-dependent quiescence contributes to tumour cell reawakening post-drug treatment. Together, our work suggests that targeting p21 function may be a viable strategy to improve outcomes for patients with *TP53*WT NSCLC.

This study identifies that *TP53*WT NSCLC cells retain an ability to enter a spontaneous, p21-dependent quiescent state in response to replication stress. We cannot conclusively link the loss of p21-dependent quiescence and subsequent increase in genomic instability to the increased cell death observed after loss of p21 in proliferating cells. However, given that, in these proliferating, unperturbed conditions, p21 levels are highest in G1 cells and that a key role of p21 appears to be in modulating the proliferation-quiescence decision during G1, we would argue that the loss of this state makes a significant contribution to the increase in spontaneous cell death after p21 loss.

At least *in vitro*, a p21-dependent quiescent state is insufficiently long to be chemoprotective. However, *in vivo*, where NSCLC cells likely proliferate much more slowly, this state could be sufficiently long to protect cells from chemotherapy and could contribute to the worse overall survival of these patients. This remains to be tested. Where more detailed patient information is available, it would be insightful to investigate if *TP53*WT p21High NSCLC patients do worse in response to chemotherapy or have higher rates of tumour relapse than p21Low patients. Whilst we did not show that pre-existing p21-dependent quiescence was chemoprotective, we did show that p21, by maintaining a 2n or 4n quiescent state post-chemotherapy, could drive tumour relapse. Many of these cells post-chemotherapy treatment are also likely to be in a p21-dependent senescent state and will never return to proliferation. However, the level of p21 is critical here. At more moderate levels of p21 expression, cells remain in a quiescent state and retain the ability to re-enter proliferative cycles (51).

Our data suggest that targeting p21 function, together with chemotherapy, could be an option to prevent the formation of a pool of p21-dependent quiescent cells post chemotherapy in *TP53*WT NSCLC. Inhibiting p21 function could prevent NSCLCs from entering, or push cells out of, p21-dependent quiescence, where they would be susceptible to treatment with chemotherapy, in a “kick and kill” approach, akin to the strategy used in HIV to eliminate latently infected cells (52,53). Previous efforts to inhibit p21, by promoting its degradation/loss, particularly in kidney cancer (54) included Sorafenib (55) and a derivative, UC2228 (56). In our hands, both failed to reduce p21 levels and did not recapitulate the known phenotypes of p21 loss. Butyrolactone (57) promotes p21 degradation but also inhibits CDKs, and LLW10 (58) promotes p21 degradation but is needed at very high concentrations (100mM). Since p21 is an intrinsically disordered protein that folds upon binding to Cyclin/CDKs, has no catalytic region to target, and occupies a large surface area on Cyclin/CDK - making it hard to target by small molecule inhibitors, peptide inhibitors may be the most viable option. Importantly, we propose a strategy to *transiently* target p21, in combination with chemotherapy. p21^-/-^ mice are developmentally normal and only develop tumours after ∼16 months (59), suggesting short-term p21 inhibition would not promote tumorigenesis. p21 loss improves lifespan in mouse models of aging, with no increase in tumorigenesis (60). Moreover, not all quiescent states are p21-dependent (37,61) plus, “reawakening” cells also requires a pro-proliferative/oncogenic stimulus, therefore we do not anticipate awakening all quiescent cells by inhibiting p21. Another option to avoid awakening quiescent cancer cells would be to push p21-dependent quiescent cancer cells into a senescent, terminally-arrested state where they could then be eradicated with senolytics (4).

Towards this latter point, our work adds to the increasing awareness of the lack of markers available to distinguish quiescent versus senescent cells. Quiescent and senescent cells share many features and thus multiparameter testing is needed to distinguish the two populations (50,62,63). The ability to detect quiescent tumour cells is important in assessing further or additional treatment options for patients to try and reduce the rates of tumour relapse. As we have mentioned and shown here, quiescence is not a single cellular state and we need a better understanding of these different states in cancer, what they mean for cancer patients and how we can better eradicate quiescent cancer cells (21). Cycles of chemotherapy aim to capture reawakening quiescent cells as they start to proliferate but the timing of these cycles must be optimised to capture reawakening quiescent cells.

Transcriptomic profiling revealed that p21High quiescent cells are enriched in transcripts encoding proteins involved in interactions with the extracellular environment, compared to proliferating cells. This is similar to what has been described in non-transformed breast epithelial cells(37) and suggests this could be a common feature of p21-dependent quiescent cells. Some of these transcripts are related to extracellular matrix (ECM) organisation, including metallopeptidases MMP2, MMP11 and ADAMTS7, ADMATS10 and ADAMTS14. In a lung cancer context, it is tempting to speculate that these quiescent cells may be capable of remodelling the ECM to initiate invasion and metastasis, as has been observed for proliferative quiescence in glioblastoma (64,65). It has been posited that invasion may even require cell cycle arrest in a “divide or conquer” strategy for cancer cells (66). It remains to be tested whether p21-dependent quiescence could contribute to invasion and metastasis in NSCLC, but if it does, then this provides another rationale for targeting p21 function in the disease.

Even after knocking out p21 in *TP53*WT NSCLC cell lines, there was often a small fraction of quiescent cells remaining, in particular in A549 and NCI-H1944 (Supplementary Figure 5D,E). This suggests that there is also a p21-independent quiescent fraction in these cells and that different quiescent states can exist in the same tumour cells. The nature of this p21-independent quiescent state remains to be determined but could be dependent on other CDK inhibitors of the Cip/Kip family – for example p27(Kip1) or p57(Kip2). It is also noteworthy how heterogeneous the p21-dependent quiescent fraction is between cell lines (Figure 2C, Supplementary Figure 3C). This does not seem to correlate with the amount of intrinsic DNA damage in these cell lines, since NCI-H460 (lowest quiescent fraction) have similar levels of endogenous damage to NCI-H1666 (second highest quiescent fraction; Supplementary Figure 4A). The balance between quiescence and proliferation depends on the amount of the CDK inhibitor, p21, and the CDK activators, D- and E-type Cyclins (21). Therefore, the fraction of cells in p21-dependent quiescence at any one time will depend on the expression levels of these proteins and the upstream pathways that influence their expression and stability.

In summary, our work reveals novel functions for p21 in *TP53*WT NSCLC patients and highlights the need for a more in-depth understanding of the different states in which cancer cells can exist to treat patients more effectively and reduce rates of tumour relapse.

## Materials and Methods

### Cell culture

A549 cells were a gift from Barbara Tanos, University of Brunel, and were STR profiled to confirm their identify. A549 and NCI-H460 (ATCC) were maintained in DMEM (Gibco 11594486). NCI-H1944 (ATCC), NCI-H1666 (ATCC), COR-L23 (Sigma) and NCI-H1650 (ATCC) were maintained in RPMI Medium 1640 (Gibco A10491-01). NCI-H1563 (ATCC), NCI-H358 (ATCC) and NCI-H1299 (ATCC) were maintained in RPMI Medium 1640 + Glutamax^TM^ (Gibco 61870-010). All media was supplemented with Penicillin/Streptomycin (P/S; Gibco, 15140-122) and 10% FBS (Sigma F9665), except for NCI-H1666 which were maintained in 5% FBS. All cell lines were maintained in incubators at 37°C and 5% CO_2_ and regularly screened for mycoplasma via in-house screening.

### Cell line generation

#### Endogenous tagging of mRuby PCNA

PCNA was labelled at the N-terminus at the endogenous locus by AAV-mediated targeting. The targeting construct pAAV-mRuby-PCNA (42) was packaged and transduced into cells, as described in (42). After four days, single cells were sorted by FACS in to 96 well plates containing 100μl 1:1 media:conditioned media. Conditioned media was filtered through 0.2 μm syringe filters to remove any cell debris. mRuby-PCNA positive clones were screened using the Operetta CLS microscope (Revvity) and positive clones expanded.

#### CDK2 reporter cell line generation

The CDK2L-GFP activity reporter, first described in (44) was transfected into A549 cells using Lipofectamine LTX reagent. One day post-transfection, cells were plated onto 15cm tissue culture plates and the next day, 0.5mg/ml G418 (Gibco) was added to select positive clones. Media and G418 was refreshed every 3-4 days and after two weeks, single-cell clones were isolated using cloning cylinders (Sigma C1059). Clones were expanded and correct localisation of the reporter was checked on the Operetta CLS microscope (Revvity).

#### p21 knockout (KO) cell lines

All p21 knockout cell lines were made as described previously (18). Briefly, cells were plated to 80% confluency one day prior to transfection with either p21KO1 or p21KO2 all-in-one nickase plasmids (18). Plasmids were transfected using Lipofectamine LTX reagent, according to the manufacturer’s instructions (Invitrogen 15338-100). Two days post-transfection, cells were single-cell sorted by FACS into 96 well plates containing 1:1 media:conditioned media. After three weeks, clones were checked for p21 expression by immunostaining. Clones with undetectable p21 expression were then expanded and checked for p21 loss by western blotting after overnight treatment with Nutlin -3 to boost p21 expression.

For genotyping, genomic DNA was extracted using the FlexiGene DNA Kit (Qiagen 51206). Nested PCR was performed using CDKN1A_KO_FWD1 5’-ATGTCAGAACCGGCT-3’, CDKN1A_KO_REV1 5’-TTAGGGCTTCCTCTT-3’, CDKN1_KO_FWD2 5’-GGATGTCCGTCAGAACCCAT-3’, CDKN1_KO_REV2 5’-GTGGGAAGGTAGAGCTTGGG-3’. PCR products were ligated into pJet1.2 (Thermo Scientific, K1232). Plasmid DNA was prepared (Qiagen, 27104) and sequenced from T7 primer by Sanger sequencing.

#### Generating p21-mVenus-AID-SMASh cell line

An mVenus-mAID-SMASh tag was introduced into the C-terminus of the *CDKN1A* gene using targeting vectors and gRNA/Cas9 cleavage, as first described in (43). Briefly, px330 p21 gRNA plasmid and pAAV-mVenus-AID-SMASh-Neo were transfected into NCI-H1944-mRuby PCNA cells using Lipofectamine LTX (Invitrogen 15338-100), according to the manufacturer’s instructions. After two days cells were plated onto 15cm dishes and allowed to attach overnight. Stable clones were selected using 0.2 mg/ml G418. After three weeks, single-cell colonies were isolated using cloning cylinders, expanded and checked by western blot and immunofluorescence.

### siRNA transfection

Cells were reverse transfected with siRNA in 384 well Phenoplates (Revvity) plates in a total volume of 20μl. All siRNAs were from Horizon Discovery: Pooled NTC D-001810-10-05, NTC D-001810-02-05, TP53 L-003329-00-0005, CDK1NA J-003471-12-0002 and used at a final concentration of 27nM. Transfection was performed using Lipofectamine RNAiMax (Invitrogen 13778150), according to manufacturer’s instructions. Briefly 40nl of Lipofectamine RNAiMax was added to 40nl 20 μM (stock concentration) siRNA in 10 μl Optimem (Gibco 31985062) and then added to cells. Transfected cells were incubated for 24-72hrs, depending on the experiment.

### Immunostaining

Cells were grown in 96 or 384 well PhenoPlates (PerkinElmer, 6055302 or 6057302, respectively). EdU was added to a final concentration of 5 μM for either 30 min (to identify S-phase cells) or 24 hr (to identify G0 cells) prior to fixation. Cells were fixed in 4% Formaldehyde/PBS for 15 min at RT, permeabilized with PBS/0.5% Triton X-100 for 15 min and blocked for 1 hr in blocking buffer (PBS/2% BSA). Primary antibodies were diluted in blocking buffer and incubated overnight at 4°C. After three washes in PBS, cells were incubated for 1 hr at RT in the dark in secondary antibodies (Invitrogen). For EdU incorporation, cells were incubated with 100 mM Tris-HCl pH 7.5, 4 mM CuS0_4_, 100 mM ascorbic acid and 5 μM sulfo-cyanine-5 azide for 30 min at RT in the dark, washed three times in PBS and counterstained with 1 μg/ml Hoechst for 10 min followed by three washes in PBS. All plate washing steps were performed on an automated 50TS microplate washer (Biotek). Plates were imaged using an Operetta CLS (Revvity) with a 20X (N.A. 0.8) objective.

Antibodies used for immunostaining were P-Rb (pSer807/811, clone D20B12, CST8516, 1:2000), 53BP1 (CST4937, 1:500), γH2AX (pSer139, CST2577, 1:1000), p21 (BD Biosciences 558430, 1:500), Goat anti mouse Alexa Fluor-488 (Invitrogen A11001, 1:1000), Goat anti mouse Alexa Fluor-568 (Invitrogen A11004, 1:1000), Goat anti mouse Alexa Fluor-647 (Invitrogen, A21235, 1:1000), Goat anti rabbit Alexa Fluor-488 (Invitrogen A11008, 1:1000), Goat anti rabbit Alexa Fluor-568 (Invitrogen A11011, 1:1000) and Goat anti rabbit Alexa Fluor-647 (Invitrogen A21245, 1:1000).

### Automated Image Analysis of Fixed Cells

All automated image analysis of Operetta CLS acquired images was performed using Harmony software (Revvity). Nuclei were segmented based on Hoescht intensity with nuclei at the edge of the field of view excluded from the analysis. Nuclear intensity of proteins was calculated as the mean intensity in the segmented region.

#### Calculating G0 Fraction

G0 cells were identified by the exclusion of P-Rb or absence of EdU staining. Briefly a nuclear to cytoplasmic (N:C) ratio was calculated by creating a four pixel width ring around the segmented nucleus, and the ring intensity calculated as a proxy for the cytoplasmic portion. N:C ratios for individual cells were calculated and plotted in Prism (Graphpad). Cells with EdU N:C ratios <1.1 were called as G0. P-Rb N:C cut-off was cell line-dependent and ranged from P-Rb < 1.4-1.8 to be called as G0.

#### Scoring 53BP1 nuclear bodies

Nuclei were segmented as above. 53BP1 nuclear bodies were segmented using the ‘Find Spots’ function in Harmony (Revvity). For untreated conditions, Method B was used with detection sensitivity 0.05 and splitting sensitivity 0.5. For aphidicolin treatments, Method C was used with radius ≤ 3.0 pixels, contrast > 0.21, uncorrected spot to region intensity > 1.2 and distance ≥ 2.0 pixels.

### Timelapse Imaging

For all live-cell imaging experiments, cells were plated in 384 well Phenoplates (Revvity) and a breathable membrane (Thermofisher) was applied to plates before imaging on the Operetta CLS (Revvity) set at 37°C and 5% CO_2_. Images were acquired at 10min intervals using a 20X (N.A. 0.8) objective. Images were exported as .tiff files and analysis was performed using NucliTrack (67) and/or FIJI. All cells analysed were present throughout the entire movie unless they died. Spontaneous death was quantified manually. mRuby-PCNA was used to define cell cycle phases as described previously (18,42).

#### Cisplatin and Gemcitabine timelapse assays

250 cells (A549-mRuby-PCNA-CDK2-GFP p21 WT or p21KO C1) and 500 cells (NCI-H1944-mRuby-PCNA) were seeded into 384 well PhenoPlates (Revvity) in 45μl of media. Cells were imaged for two days before 5 μl of Cisplatin (Sigma P4394) or Gemcitabine (APExBIO A8437-APE) was added to 16μM and 5μM, respectively, before imaging for a further three days.

#### Spontaneous Death Experiments

250 A549-mRuby-PCNA-CDK2-GFP p21 WT or 500 NCI-H1944-mRuby-PCNA cells were plated in 20μl of media in duplicate plates. p21 siRNA was performed as described. One plate was fixed after 6hrs and, at the time, the second plate was imaged by timelapse imaging. After three days this second plate was fixed then both plates were immunostained for p21 to ensure knockdown at the start and end of the experiment.

### Growth Curves

Cells were plated at a density of either 1000 cells (A549 and NCI-H460 and derivatives) or 2000 cells (NCI-H1944 and derivatives) in triplicate in 96 well tissue culture plates. Brightfield images were taken every 4 hrs for 14 days using the 10x (N.A. 0.3) objective and percent confluency calculated on the CellCyteX (Echo) live imaging system. To calculate confluency a mask was created using the default settings on the CellCyteX (brightness 0%, contrast 0%, contrast sensitivity 50 a.u., smoothing a.u. and filled hole size 100µm^2^). Results were plotted in Prism (Graphpad) with standard deviations.

### RNA-seq

Five million cells were plated on to 600cm^2^ plates. Cells were left for a total of four days with media changed on day two. On collection for RNA-seq experiments 2000 cells were plated in 384 well Phenoplates and left to attach for 6 hrs before fixing and immunostaining, for quality control checks. 5 μM EdU was added 1 hr prior to fixation. Cells were immunostained for proliferation/quiescence markers, namely EdU incorporation, P-Rb and p21.

Total RNA was extracted using a RNeasy MinElute Cleanup Kit (Qiagen 74204) according to manufacturer’s instructions. Quality and concentration was assessed using the Agilent 2100 Bioanalyser RNA 6000 Nano assay.

PolyA enrichment was performed using NEBNext^®^ *Poly(A) mRNA Magnetic Isolation module,* and NGS RNASeq libraries made using the NEBNext® Ultra™ II Directional RNA Library Prep Kit for Illumina® according to manufacturer’s instructions.

Library size and adaptor contamination was assessed using the Agilent 2100 Bioanalyser High Sensitivity DNA assay, and concentrations measured with the Qubit dsDNA High Sensitivity assay.

For each sample, a minimum of 50 million Paired End 60bp reads were generated on an Illumina NextSeq2000 with unique dual 8bp indexing.

### Western Blotting

Whole cell lysates were prepared following aspiration of media from culture plates, followed by washing with PBS on ice. Lysates were collected in 1X Novex Tris-Glycine-SDS sample buffer (Novex, LC2676) supplemented with 1x phosphatase inhibitors (ThermoScientific, 1862495), 1x protease inhibitors (ThermoScientific, 78429) and 1 mM DTT (BioUltra, 43816-10ML). Samples were incubated at 95 °C for 10 min then centrifuged at 14,000 xg for 1 min before loading on Novex 4-20% Tris-Glycine gel (Invitrogen, XP04205). After transfer to PVDF-FL (Merck Life Sciences, IPFL00010), membranes were blocked in blocking buffer (TBS, 5% milk, 10% glycerol, Tween 0.1%) for 1hr at RT and incubated in primary antibodies diluted in blocking buffer, overnight at 4°C in. Membranes were washed three times in TBS/0.05% TritonX-100 before incubating for 1hr at RT with HRP-conjugated secondaries, diluted in blocking buffer. Membranes were washed three times in TBS/0.05% TritonX-100 and developed using Clarity Western ECL Substrate (Bio-Rad, 1705061). Blots were imaged on an Amersham Imager 680. Where needed to boos p21 signal, Nutlin-3 (Sigma N6287) was added to a final concentration of 5 μM (A549 and NCI-H460) or 10 μM (NCI-H1944) for 24 hr before cell lysates were prepared.

Antibodies used for western blotting were p21 (BD Biosciences 558430, 1:500), Vinculin (E1E9V, CST13901, 1:1000), Anti mouse-HRP (CST 7076P2, 1:1000) and Anti rabbit-HRP (CST 7074P2, 1:1000).

### Relapse Assay

Cells were plated in triplicate wells on 96 well PhenoPlates in the following numbers and allowed to attach overnight: 4,500 (NCI-H460, NCI-H460 p21KO C11, A549-mRuby-PCNA-CDK2L-GFP and A549-mRuby-PCNA-CDK2L-GFP p21KO C1),

10,000 (NCI-H1944) and 15,000 (NCI-H1944 p21KO C59). The next day, cells were imaged on the CellCyteX (Echo) for 24 hr before addition of cisplatin, etoposide or gemcitabine at 16μM, 5μM and 5μM, respectively. After three days, plates were washed with media and imaging was continued for a total of 28 days. Percentage confluency was calculated using CellCyteX software as a proxy for growth. After 28 days cells were stained with Hoechst and imaged on the Operetta CLS (Revvity) using a 20x (N.A. 0.8) objective as a second way to measure assay endpoint. These data were analysed in ImageJ by calculating the percentage of Hoechst coverage/well. Briefly images were converted to 16-bit and sliding scale thresholds altered to cover all stained areas. In all cases, the upper threshold was kept at the highest value. The lower threshold varied slightly between cell lines. Area was calculated using the “measure” function.

### FACS assay for G1 and G2 regrowth

Two million NCI-H1944 cells were plated and allowed to attach for 24 hrs. Gemcitabine was added to a final concentration of 5 μM and cells incubated in drug for 72 hrs. Cells were sorted into G1 and G2 fractions using a BD FACSAria cell sorter with BDFACSDiva software v9.4 and cells re-plated at 1000 cells/96 well in triplicate. After three weeks cells were stained with Hoescht. Plates were imaged on an Operetta CLS using a 20x (N.A. 0.8) objective and percentage Hoescht coverage per well was calculated using ImageJ, as described above.

### Ebvacociclib Assay

2000 NCI-H1944 cells were plated in 90 μl in triplicate on 96 well Phenoplates (Revvity) and left for 48 hrs. DMSO or ebvaciclib (PF06873600, Caltag Medsystems Ltd, TAR-T8563-1mg) was added to cells for 24 hrs. Plates were washed three times and fresh media added containing 5 μM EdU to monitor the ability of cells to proliferate. To this, 10 μl of a 10X solution containing either drug (cisplatin, etoposide or gemcitabine) and ebvaciclib or controls was added to cells for three days. Cells fixed and stained for Hoechst and EdU, as described previously. Plates were imaged on the Operetta CLS (Revvity) using a 20x (N.A. 0.8) objective.

### Drug dose curves

Cells were plated in triplicate wells in 45 μl of media on 384 well PhenoPlates in the following numbers and allowed to attach overnight: 500 cells (A549-mRuby-PCNA-CDK2-GFP p21 WT or p21KO C1/C6 and NCI-H460 or NCI-H460 p21KO C11) or 1000 cells (NCI-H1944 and NCI-H1944 p21KO C59). A 10X 1:5 serial dilution of either cisplatin, etoposide or gemcitabine was prepared in growth media and 5 μl added to each well. Final concentrations for all assays were 100, 20, 4, 0.8, 0.16, 0.032, 0.0064, 0.0013 and 0.0003 μM. Plates were incubated for three days. Cells were fixed and stained for Hoechst and survival calculated as a percentage of vehicle. IC_50_ values were calculated using Prism (Graphpad).

#### Immunohistochemistry on patient tumours

Tumours were fixed in 10% neutral buffered formalin, embedded in paraffin and 3 μm sections prepared. Tumour sections were demarcated by pathologist Saral Desai.

Immunohistochemistry was carried out on sections after deparaffinization, re-hydration in a descending Ethanol series and antigen retrieval in Tris-EDTA pH 9.0 antigen retrieval buffer (Abcam ab93864) at 95°C for 30min. Sections were blocked (PBS, 10% rabbit serum, 1% BSA, 0.1% triton x-100, 0.01% tween, 0.2% coldwater fish gelatin) for 2 hrs at RT. Primary antibodies were incubated overnight at 4°C in blocking buffer with 5% rabbit serum. Sections were washed and incubated with secondary antibodies at RT, in the dark for 1 hr. The TruView Autofluorescence Quenching Kit (Vector Laboratories SP-8400) was used after washing to decrease background followed by counterstaining with DAPI.

Antibodies used for immunohistochemistry were Ki67 (Abcam, ab15580, 1:50), p21 (BD Biosciences, 558430, 1:50), Goat anti rabbit Alexa Fluor-488 (Invitrogen, A11008, 1:500) and Goat anti rabbit Alexa Fluor-647 (Invitrogen, A21245, 1:500).

Sections were imaged on a Zeiss Axio Scan Z1 and data batch analysed using QuPath (68). Briefly, following nuclei detection using Stardist (69) with a pretrained model followed by cell expansion, the mean nuclear to cytoplasm intensity ratio was calculated for each respective channel, before classification of cell type using a composite object classifier.

### Analysis of NSCLC human cohorts

FPKM normalised RNA-seq data for NSCLC patients (lung adenocarcinoma and lung squamous cell carcinoma) were downloaded from the Human Protein Atlas (https://www.proteinatlas.org/about/download). Protein RPPA levels and clinical information for the same individuals were obtained from cBioPortal (https://www.cbioportal.org/).

Groups of p21 high and p21 low-expressing tumours were defined using the 75% and 25% quartiles of the p21 (gene and protein) expression distribution, respectively. Survival analysis and Kaplan-Meier curves comparing the two groups were generated using the *survival*, *survminer* and *ggpubr* R packages, respectively.

## Data availability

The data generated in this study are publicly available in Gene Expression Omnibus (GEO) at GSE266945.

## Acknowledgements

We thank MRC-LMS flow cytometry, genomics, bioinformatics and microscopy facilities for their technical support throughout this project. ARB and SJC were supported by a CRUK Career Development Fellowship to ARB (C63833/A25729), and work in the lab was also supported by MRC-LMS core funding (MC-A658-5TY60). MS was supported by a UKRI Future Leaders Fellowship (MR/T042184/1). Work in MS’s lab was supported by a BBSRC equipment grant (BB/R01356X/1) and a Wellcome Institutional Strategic Support Fund (204841/Z/16/Z). LVF and POP are supported by MRC-LMS core funding (MC-A654-5QC70) and work in the lab is also supported by a CRUK Career Establishment Award to LVF (RCCCEA-Nov21\100001). FAH was supported by a studentship from the EPSRC Centre for Mathematics in Precision Healthcare (EP/N014529/1). PT is supported by a UKRI Future Leaders Fellowship (MR/T018429/1). Human samples used in this research project were obtained from the Imperial College Healthcare Tissue Bank (ICHTB). ICHTB is supported by the National Institute for Health Research (NIHR) Biomedical Research Centre based at Imperial College Healthcare NHS Trust and Imperial College London. ICHTB is approved by Wales REC3 to release human material for research (17/WA/0161), and the samples for this project (R20007-1A) were issued from sub-collection reference number HIS_SD_18_025.

