## Supplementary Figures and Legends for "Pro-survival roles for p21(Cip1/Waf1) in Non-Small Cell Lung Cancer"

### Supplementary Information

### Supplementary Figure Legends

**Supplementary Figure 1.** TCGA analyses showing overall survival probability separated by *TP53* status and level of p21 protein expression.

**Supplementary Figure 2. A.** Representative graphs show that *TP53*WT NSCLC cell lines retain a functional p53-p21 response to treatment with the Mdm2 inhibitor, Nutlin-3. Cells were treated for 24h with 5 mM Nutlin-3 and then fixed and stained for p21. **B.** Representative graphs show that *TP53*WT NSCLC cell lines retain p53-dependent expression of p21 in an unperturbed condition. Red curves indicate levels of p21 expression, quantified by immunostaining, after p53 siRNA treatment of cell lines. NTC is non-targeting control siRNA.

**Supplementary Figure 3. A.** Schematic shows how phosphorylated S807/811 pRb (P-Rb) can be used to quantify proliferating versus quiescent (G0) cells. **B.** Sample images from quantitative single-cell imaging to measure pS807/811 pRb levels and p21 expression in NSCLC cell lines. White arrows indicate cells that are in quiescence with high p21 expression. Scale bar 50 mm. p21 is green and pS807/811 Rb is red in merged image. **C.** Graph shows quantification of percentage of P-Rb negative (G0) cells across cell lines. Data are plotted as superplots, n=3, 4 technical replicates per experiment. Grey dot represents the mean. *TP53*WT cell lines are shown in red, psi-p53 mutant is shown in purple and *TP53*mutant cell lines in blue. **D.** Graph shows correlation between fraction of quiescent cells quantified by EdU and P-Rb assay. Each dot represents the mean of n=3. **E.** Graph showing depletion of p21 by p21 targeting siRNA, quantified by immunostaining. Cells treated with non-targeting control (NTC) siRNA are shown in black. Cells treated with p21-targeting siRNA are shown in red. **F.** Graph shows the effect of p21 depletion by siRNA on the fraction of quiescent cells within each cell line, measured using the P-Rb assay. Data are plotted as superplots, n=3, 4 technical replicates per experiment. NTC values are shown in black, p21 siRNA in red. **G.** Western blot showing homozygous tagging of endogenous p21 with mVenus-AID-SMASh tag. Nutlin was added for 48h to boost p21 levels. Vinculin was used as a loading control. Parent refers to NCI-H1944 mRuby-PCNA cell line. Tagged refers to NCI-H1944 mRuby-PCNA p21-mVenus-AID-SMASh cell line. **H.** Growth curves of tagged and untagged NCI-H1944 cells to show that tagging does not perturb cell proliferation. **I.** Still images from NCI-H1944 mRuby-PCNA p21-mVenus expressing cells taken from timelapse imaging. Scale bar 10 mm. **J.** Two example graphs of mRuby-PCNA and p21-mVenus quantification from nuclei of segmented and tracked NCI-H1944 mRuby-PCNA p21-mVenus cells. mRuby-PCNA is used to define cell cycle transitions. Data is plotted around time of mitotic exit. **K.** Still images of A549 cells expressing the CDK2 activity reporter (CDK2L-GFP; upper panels) and mRuby-PCNA (lower panels) showing single-cell heterogeneity in quiescence entry in genetically identical cell populations. Time is from mitotic exit of indicated cells. Both sister cells enter a period of low CDK2 activity post-mitosis. The white arrow indicates a cell that enters S-phase at 770 mins (12.8h) post-mitosis. The yellow arrow indicates a cell that remains in quiescence after 1200 mins (20h) post-mitosis. Scale bar is 20 □m. **L.** Example FACS plot showing the gates used to sort $p21^{High}$ $PCNA^{Low}$ (quiescent) and $p21^{Low}$ $PCNA^{High}$ (proliferating) NCI-H1944 p21-mVenus cells. **M.** Graphs showing validation of cell states by immunostaining post-FACS sort. Briefly, sorted cells from the two populations (p21-High and p21-Low), and cells FACS sorted from the total population (Total) were plated into 384 well CellCarrier plates and fixed and immunostained 6h post-sorting. **N.** PCA plot showing separation of p21High and p21Low clusters in PC1.

**Supplementary Figure 4. A.** Graphs show the percentage of p21-High (upper panels) or EdU negative (lower panels) quiescent cells with a 53BP1 nuclear body, compared to proliferating cells. Data are representative of n=3. **B.** Representative images of A549 cells treated with low-dose (0.3 mM) aphidicolin for 72h. Hoechst is shown in blue, EdU in red, P-Rb in orange and p21 in green in merged images. Scale bar 100 mm. **C.** Representative images of A549 cells treated with low-dose (0.3 mM) aphidicolin for 72h. Hoechst is shown in blue, EdU in red, p21 in green and 53BP1 in orange in merged images. Scale bar 100 mm. **D.** Quantification of percentage of cells with at least one 53BP1 nuclear body after aphidicolin treatment. **E.** Quantification of p21 levels after aphidicolin treatment in single-cells, representative of n=3. Mean +/- std is shown in black.

**Supplementary Figure 5. A.** Graphs of cell cycle progression for A549 mRuby-PCNA cells transfected with NTC or p21-targeting siRNA. 34 cells for NTCsi and 32 cells for p21si. **B.** Western blots to verify p21KO cell lines. 10 mM Nutlin-3 was added (where indicated) for 48h to boost p21 signal. Vinculin is used as a loading control. **C.** Growth curves of p21WT and p21KO cell lines. Data are shown as mean +/- stdev for three technical replicates. **D.** Percentage of EdU negative cells in p21WT versus p21KO. Cells were labelled with EdU for 24h before fixing and staining. Data are presented as superplots, n=3, 4 technical repeats per experiment. Black line represents mean +/- stdev. **E.** Percentage of P-Rb negative cells in p21WT versus p21KO. Data are presented as superplots, n=3, 4 technical repeats per experiment. Black line represents mean +/- stdev. **F.** Quantification of gH2AX levels across all cell cycle phases in p21WT and p21KO clones across three NSCLC cell lines. Data are representative of n=3.

**Supplementary Figure 6. A.** Dose curves of A549 (red), NCI-H1944 (purple) and NCI-H460 (blue) cells treated with etoposide (eto), gemcitabine (gem) or cisplatin (cis) for three days. Grey boxes indicate surviving fraction of cells, even at the highest doses. Data are representative of n=1. **B.** Correlation between fraction of cells surviving gemcitabine (left) or cisplatin (right) treatment and quiescent fraction in that cell line. Cells were treated for three days with 5 mM gemcitabine or 16 mM cisplatin. Mean +/- std for each repeat plotted and n=2 for each cell line. **C.** p21WT and p21KO cell lines were treated for three days with 5 mM of gemcitabine (left) or 16 mM of cisplatin (right) and surviving fraction was calculated. Data are plotted as superplots, n=3 biological repeats and four technical repeats per experiment. Black line represents the mean. One-way ANOVA was used to calculate statistical significance. **** = p<0.0001, ** = p<0.01, ns = not significant. **D.** Cell cycle phase plots for NCI-H1944 mRuby-PCNA cells treated with gemcitabine or cisplatin for three days. Cells were imaged for two days before drug addition (marked by vertical black lines at 2880 mins). Tables underneath show the percentage of cells that started in each fate upon drug addition (G1, S or G2) and their end fate (G1, S, G2 or death (D)). **E.** Graphs showing percentage of proliferating NCI-H1944 cells (EdU positive) after addition of CDK2/4/6i and chemotherapy agent. EdU was added at the same time as chemotherapy i.e. one day after CDK2/4/6i addition. Data are plotted as superplots, n=3 biological replicates with three technical replicates per repeat. Black line represents the mean.

**Supplementary Figure 7. A.** Representative images pf NCI-H1944 p21WT (upper panels) and p21KO (lower panels) cells 24 days post-washout of three day vehicle or gemcitabine treatment. Hoechst is in blue and EdU in red in merged images. EdU was added for the last 24h before fixing. Scale bar is 1mm. **B.** Quantification of percentage of well covered by A549 nuclei at the end of the relapse assay (28 days). Mean +/- stdev of n=3 are shown and individual wells are plotted for each technical repeat. **C.** Quantification of percentage of well covered by NCI-H460 nuclei at the end of the relapse assay (28 days). Mean +/-stdev of n=3 are shown and individual wells are plotted for each technical repeat. **D.** Phase contrast images from CellCyteX long-term imaging to highlight different morphologies of cells that regrow after gemcitabine treatment. Two main morphologies are observed 3.5 weeks after drug washout – small, tightly-packed cells (bounded by black dashed line, top left) and larger, more spread cells (remainder of field of view). Scale bar is 100mm. Time is in hours. **E.** FACS plots showing gates used to sort gemcitabine-treated G1 and G2 cells. Cells were labelled with Hoechst 33258 before sorting.

**Table S1 –** GO terms of transcripts significantly decreased in p21High cells compared to p21Low cells.

**Table S2 -** GO terms of transcripts significantly increased in p21High cells compared to p21Low cells.

**Table S3 –** Transcripts per million (TPM) of all genes across the human genome in p21High and p21Low conditions.

**Movie S1.**
NCI-H1944 cells expressing mRuby-PCNA (left panel) and p21-mVenus (right panel).

**Movie S2.**
A549 cells expressing mRuby-PCNA (left panel) and CDK2L-GFP (right panel).

**Movie S3.**
NCI-H1944 p21WT cells treated with vehicle. Vehicle was added at 1d for a further 3d, then washed out and cells imaged for 24 days. Time is in hrs.

**Movie S4.**
NCI-H1944 p21WT cells treated with gemcitabine. Gemcitabine was added at 1d for a further 3d, then washed out and cells imaged for 24 days. Time is in hrs.

**Movie S5.**
NCI-H1944 p21KO cells treated with vehicle. Vehicle was added at 1d for a further 3d, then washed out and cells imaged for 24 days. Time is in hrs.

**Movie S6.**
NCI-H1944 p21KO cells treated with gemcitabine. Gemcitabine was added at 1d for a further 3d, then washed out and cells imaged for 24 days. Time is in hrs.

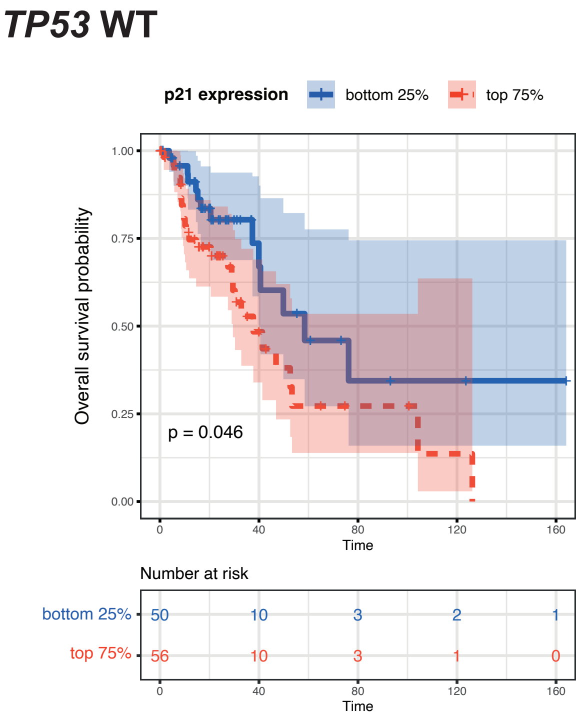


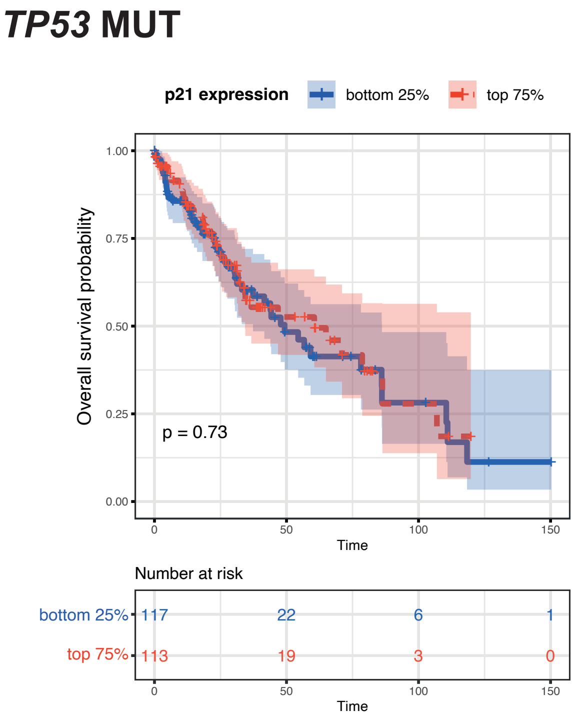


**Supplementary Figure 1**

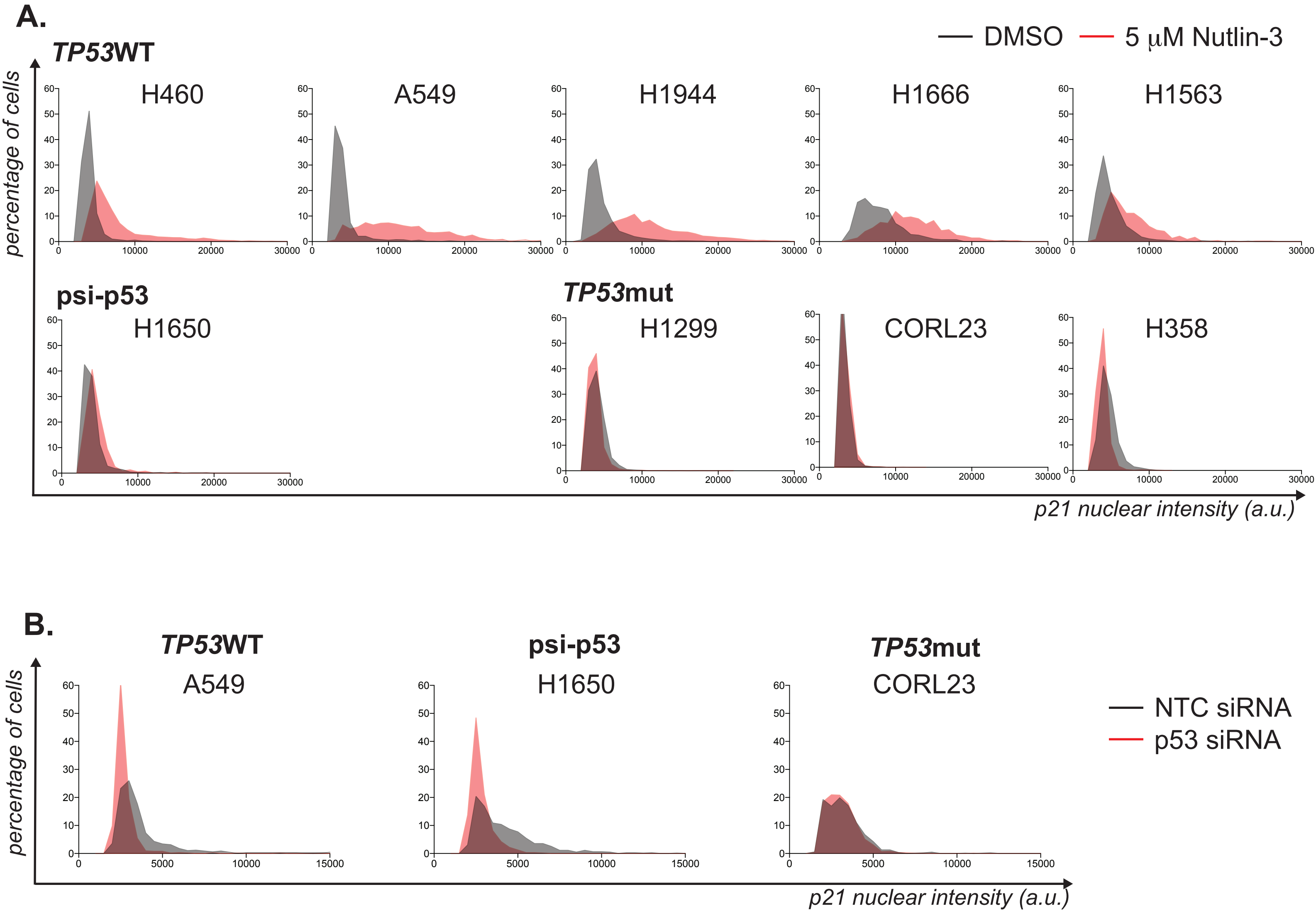


**Supplementary Figure 2**

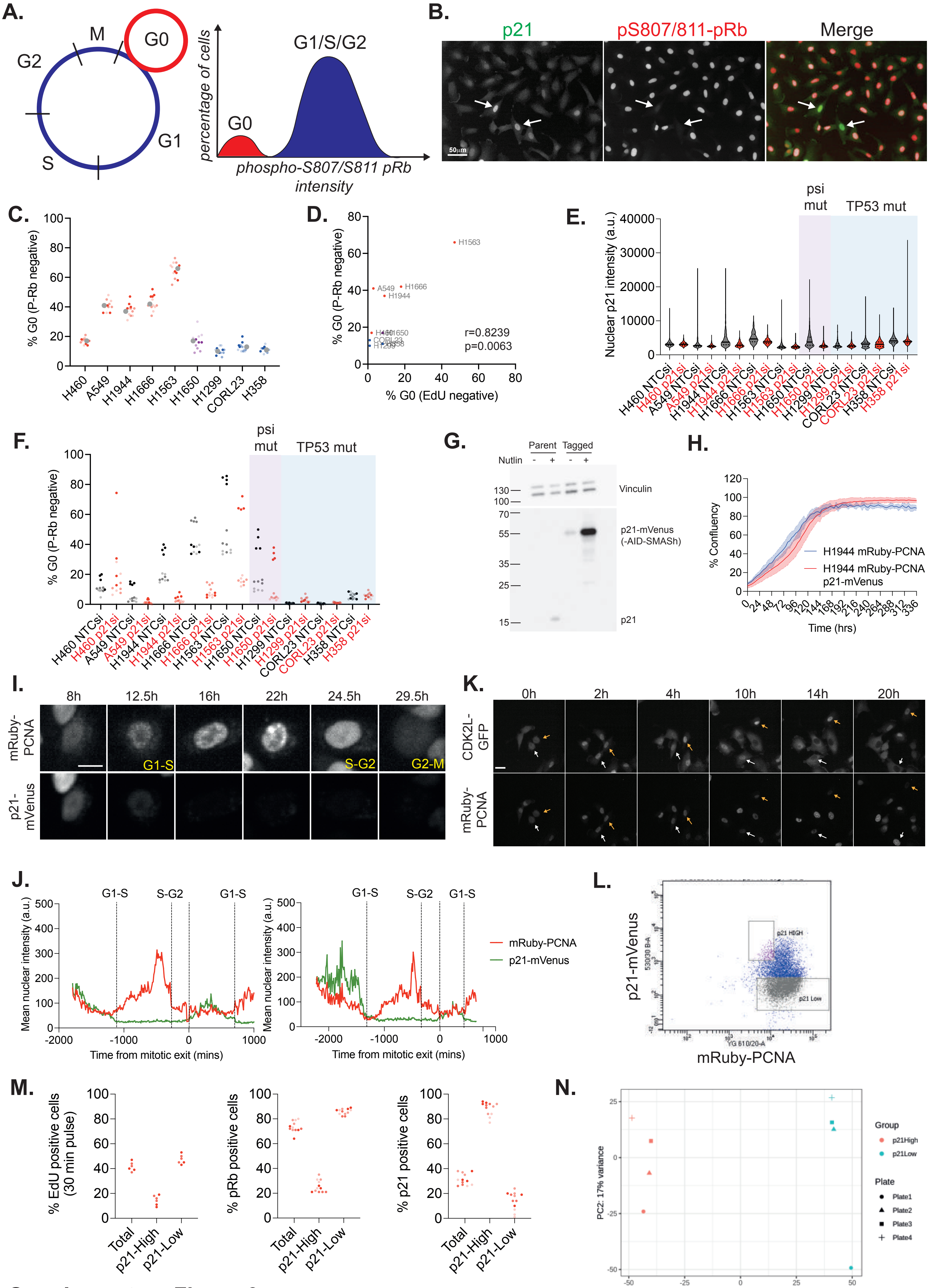


**Supplementary Figure 3**

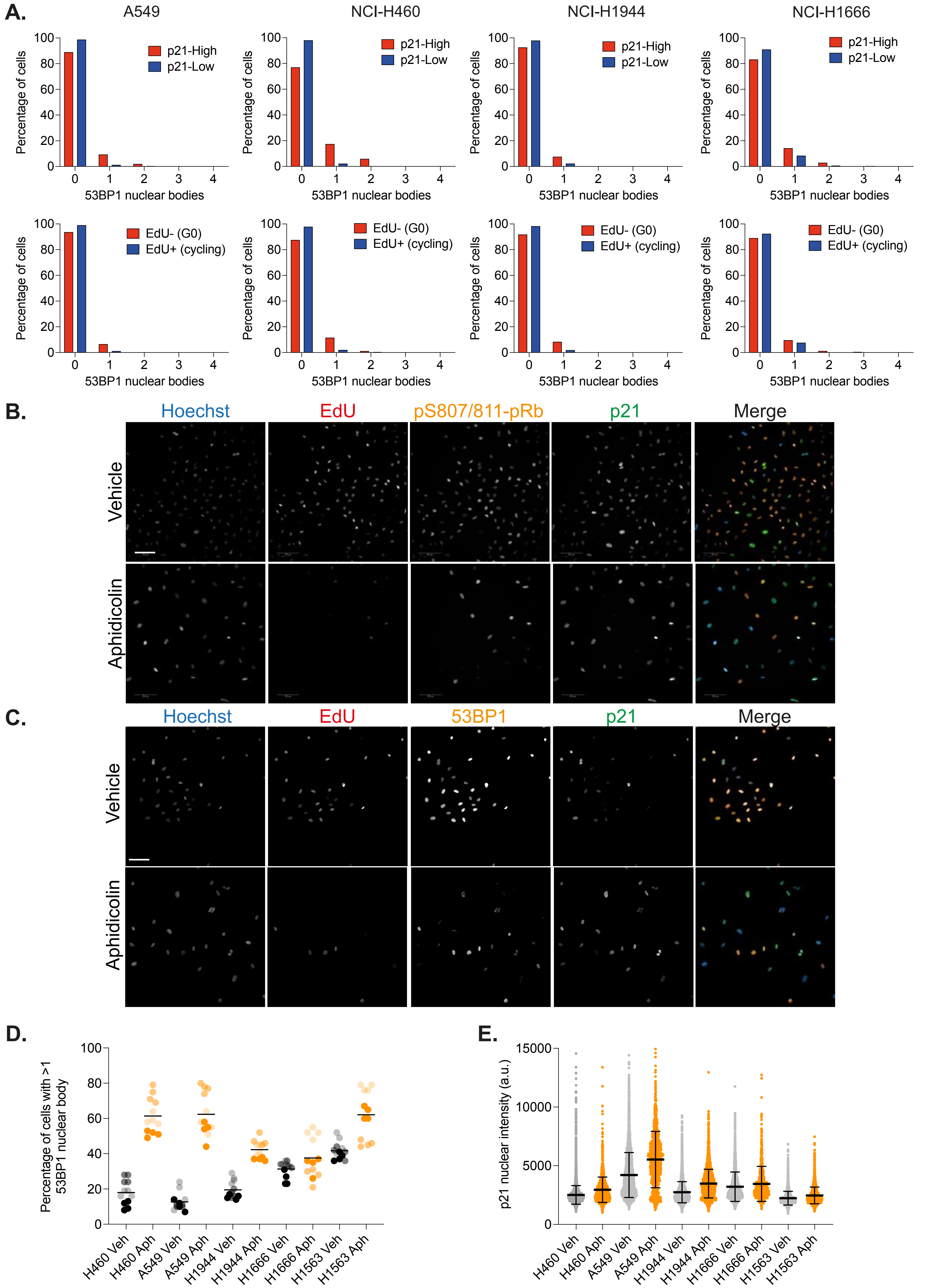


**Supplementary Figure 4**

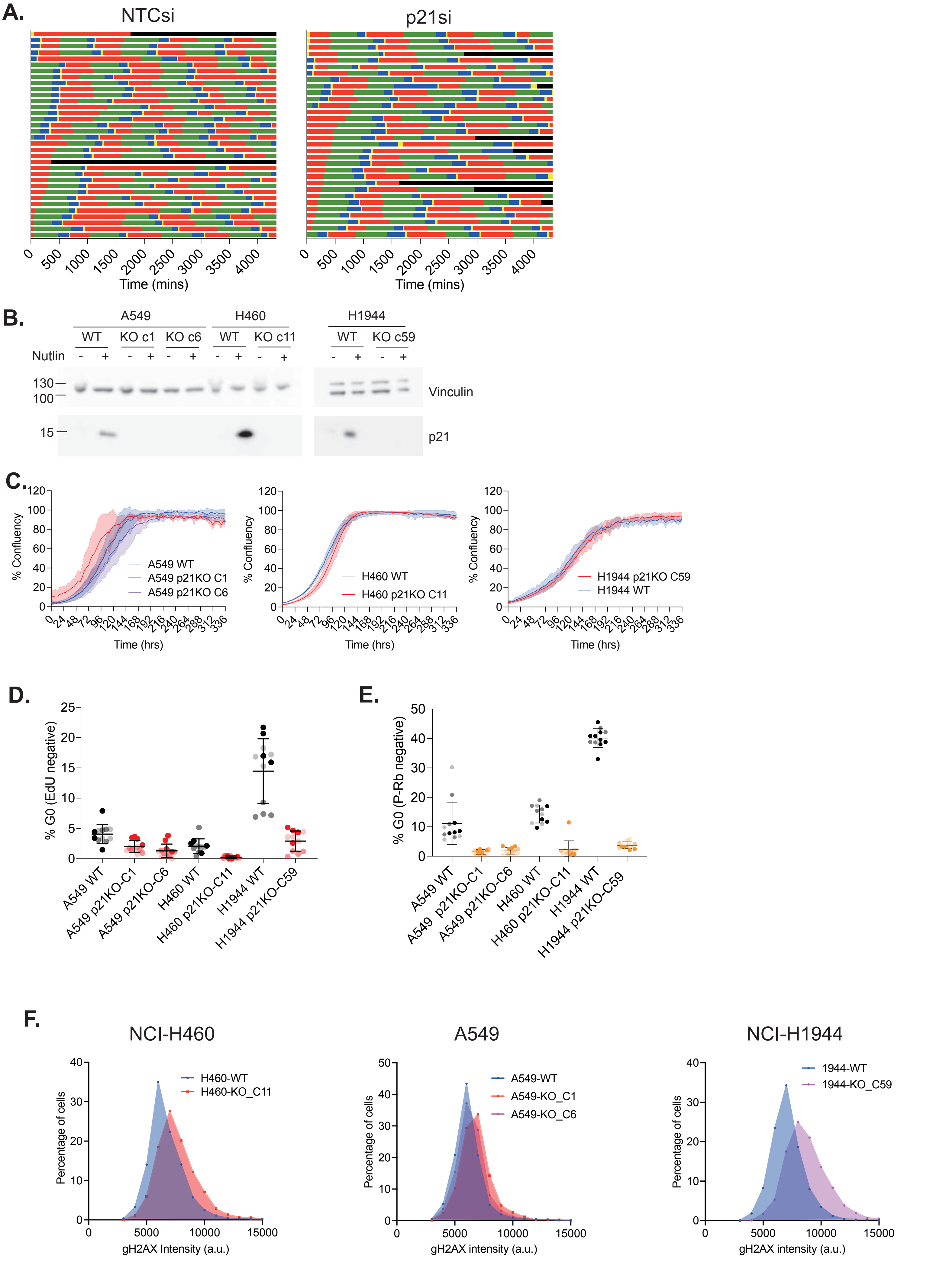


**Supplementary Figure 5**

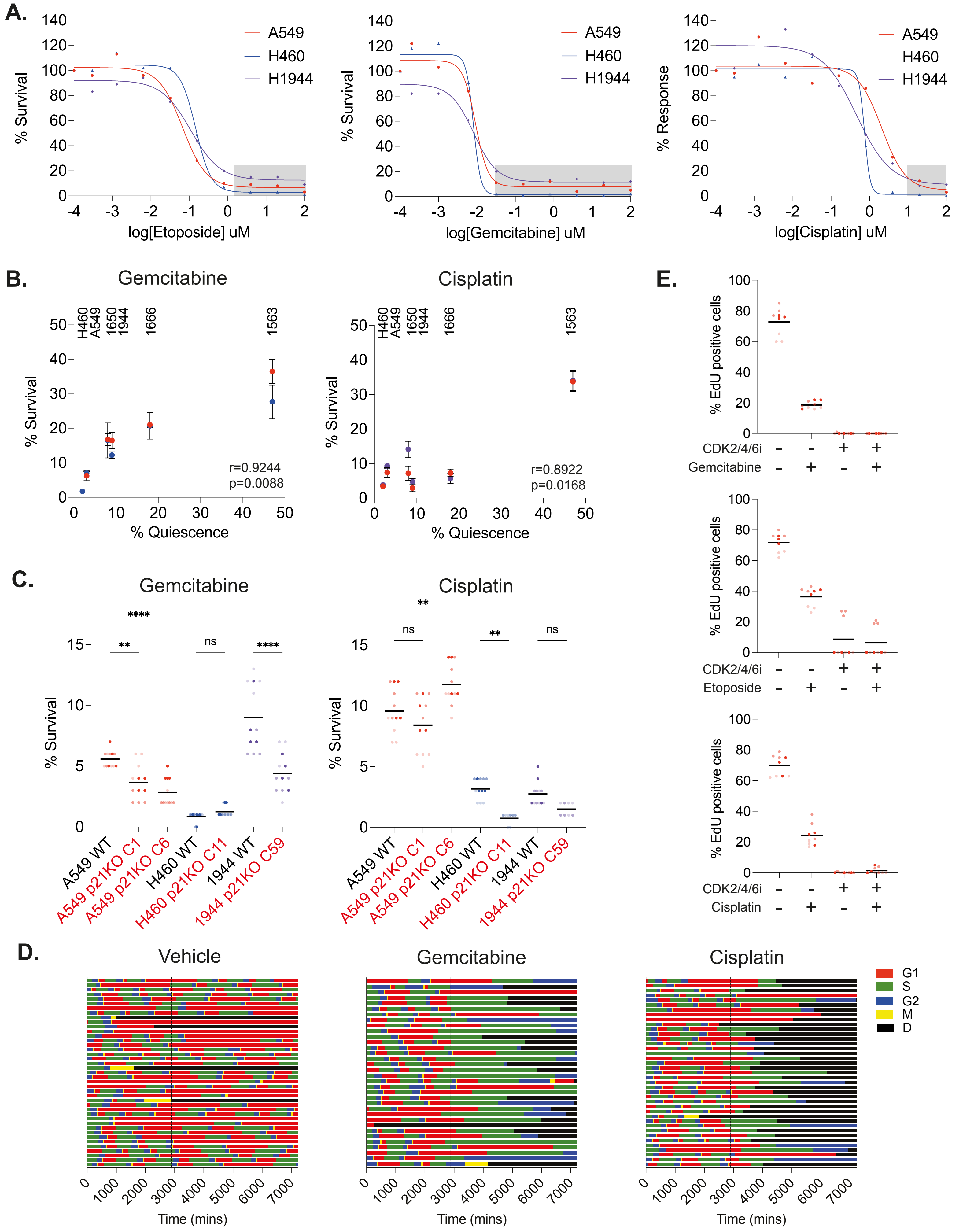


**Gemcitabine**

| End fate \ Start phase | G1 | S | G2 |
|---|---|---|---|
| G1 | 10 | 0 | 0 |
| S | 10 | 15.8 | 25 |
| G2 | 30 | 26.3 | 25 |
| D | 50 | 57.9 | 50 |

**Cisplatin**

| End fate \ Start phase | G1 | S | G2 |
|---|---|---|---|
| G1 | 0 | 0 | 20 |
| S | 0 | 0 | 0 |
| G2 | 13 | 11.1 | 0 |
| D | 87 | 88.9 | 80 |

**Supplementary Figure 6**

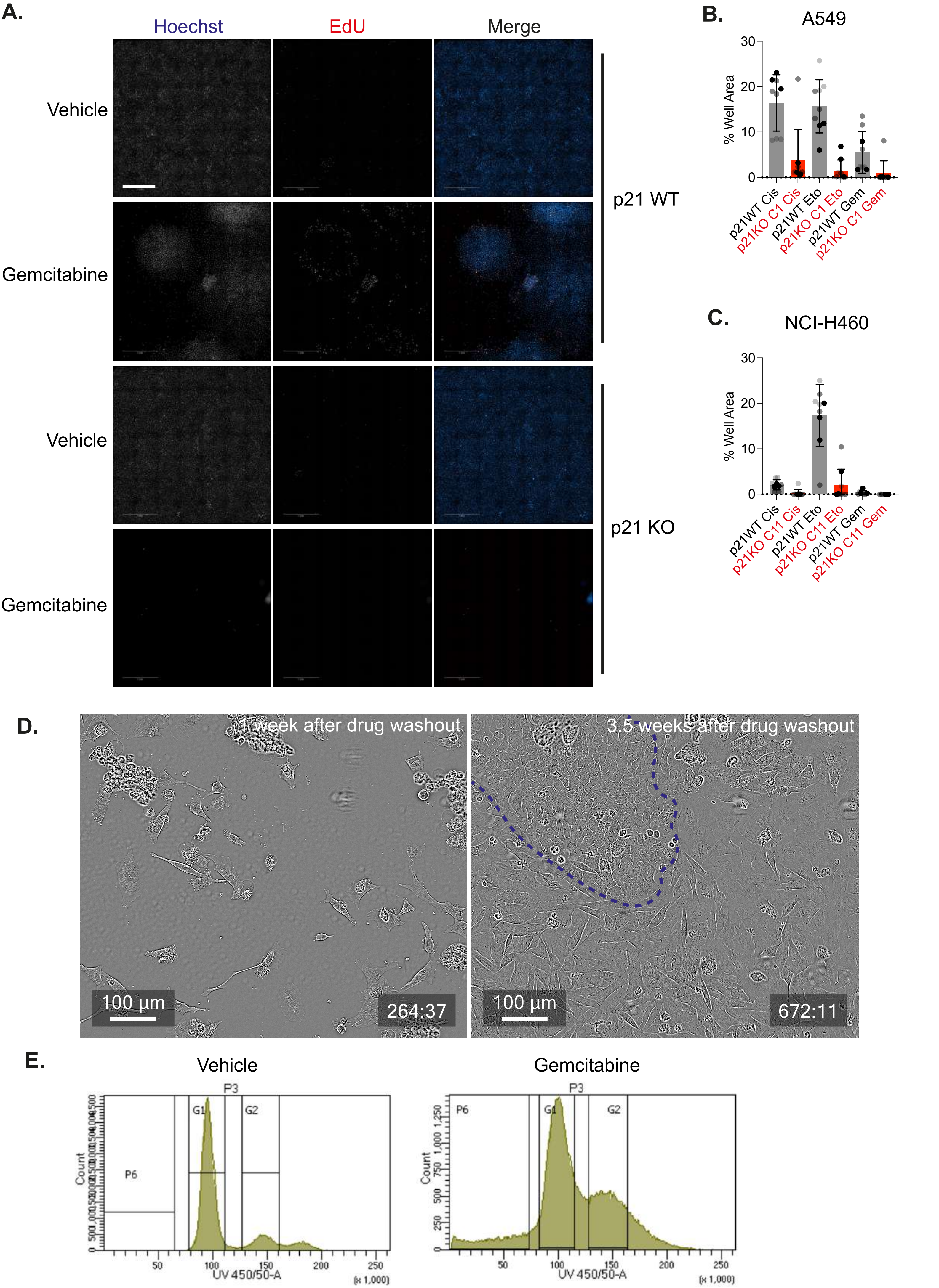


**Supplementary Figure 7**
